# Mechanism of hERG inhibition by gating-modifier toxin, APETx1, deduced by functional characterization

**DOI:** 10.1101/2020.07.22.215491

**Authors:** Kazuki Matsumura, Takushi Shimomura, Yoshihiro Kubo, Takayuki Oka, Naohiro Kobayashi, Shunsuke Imai, Naomi Yanase, Madoka Akimoto, Masahiro Fukuda, Mariko Yokogawa, Kazuyoshi Ikeda, Jun-ichi Kurita, Yoshifumi Nishimura, Ichio Shimada, Masanori Osawa

**Affiliations:** Graduate School of Pharmaceutical Sciences, Keio University, Shibakoen, Minato-ku, Tokyo, 105-8512, Japan; Division of Biophysics and Neurobiology, Department of Molecular and Cellular Physiology, National Institute for Physiological Sciences, Myodaiji, Okazaki-shi, Aichi, 444-8585, Japan; Nanion Technologies Japan K.K., Tokyo Laboratory, Wakamatsu-cho, Shinjuku-ku, Tokyo 162-0056, Japan; Institute for Protein Research, Osaka University, Yamadaoka, Suita-shi, Osaka, 565-0871, Japan; NMR Science and Development Division, RSC, RIKEN, Suehiro-cho, Tsurumi-ku, Yokohama-shi, Kanagawa 230-0045, Japan; Graduate School of Pharmaceutical Sciences, The University of Tokyo, Hongo, Bunkyo-ku, Tokyo 113-0033, Japan; Graduate School of Medical Life Science, Yokohama City University, Suehiro-cho, Tsurumi-ku, Yokohama-shi, Kanagawa, 230-0045, Japan

## Abstract

Human *ether*-*à*-*go*-*go*-related gene potassium channel 1 (hERG) is a voltage-gated potassium channel, the voltage-sensing domain (VSD) of which is targeted by a gating-modifier toxin, APETx1. Although it is known that APETx1 inhibits hERG by stabilizing the resting state, it remains unclear where and how APETx1 interacts with the VSD in the resting state. Here, we prepared a recombinant APETx1, which is structurally and functionally equivalent to the natural product. Electrophysiological analyses using wild type and mutants of APETx1 and hERG revealed that their hydrophobic residues, in addition to a previously reported acidic hERG residue, play key roles in the inhibition of hERG by APETx1. Docking models of the APETx1-VSD complex that satisfy the results of mutational analysis suggest a molecular recognition mode between APETx1 and the resting state of hERG; this would provide a structural basis for designing ligands that control hERG function by binding to the VSD.

## Introduction

Human *ether*-*à*-*go*-*go*-related gene potassium channel 1 (hERG; K_V_11.1) is a voltage-gated potassium channel (K_V_ channel) expressed in human cardiomyocytes, as well as brain and cancer cells (*Sanguinetti and Tristani-Firouzi, 2006; Vandenberg et al., 2012; Warmke and Ganetzky, 1994*). hERG conducts potassium ions (K^+^) across the cell membrane upon depolarization, thereby contributing to the repolarization of the action potential (*Sanguinetti and Tristani-Firouzi, 2006; Vandenberg et al., 2012*). This function is necessary for a normal heartbeat, as demonstrated by the fact that some hERG inhibitors cause lethal arrhythmia accompanied by long QT syndrome (*Curran et al., 1995; Roden, 2004; Sanguinetti et al., 1995; Sanguinetti and Tristani-Firouzi, 2006*). Recently, it has been reported that variations in the gene encoding hERG are associated with schizophrenia (*Apud et al., 2012; Atalar et al., 2010; Huffaker et al., 2009*), and that alterations in hERG expression and function are observed in various types of cancer cells and are involved in carcinogenic processes (*He et al., 2020; Jehle et al., 2011; Rao et al., 2015*). Non-arrhythmogenic hERG inhibitors, which block hERG without inducing arrhythmia, can improve the survival rate among glioblastoma patients showing high hERG expression (*Pointer et al., 2017*). These clinical results demonstrate the therapeutic potential of the specific ligands controlling hERG function; thus, it is of great medical importance to determine the structural mechanisms underlying the interactions between hERG and its specific ligands (*Arcangeli and Becchetti, 2017; He et al., 2020; Vandenberg et al., 2012*).

hERG is a tetrameric channel in which each subunit comprises six transmembrane segments (S1-S6) and N- and C-terminal cytoplasmic domains (*Ben-Bassat et al., 2020; Li et al., 2016; Morais Cabral et al., 1998; Wang and MacKinnon, 2017; Warmke and Ganetzky, 1994*). In the tetrameric architecture of K_V_ channels, the S5 and S6 segments form a pore domain (PD) with a K^+^-selective filter and an activation gate at the center of the tetramer, and the S1-S4 segments of each subunit form a voltage-sensing domain (VSD) at the four peripheries of the PD (*Chen et al., 2010; Long, 2005; Sun and MacKinnon, 2017; Sun and MacKinnon, 2020; Wang and MacKinnon, 2017; Whicher and MacKinnon, 2016*). The voltage-dependent conformational changes of the VSD regulate the opening and closing of the gate in the PD (*Bezanilla, 2008; Börjesson and Elinder, 2008; Swartz, 2008*). Upon membrane depolarization, the VSD undergoes a conformational change from “S4-down” to “S4-up,” in which S4 moves from the intracellular side to the extracellular side roughly perpendicular to the membrane plane (*Bezanilla, 2008; Börjesson and Elinder, 2008; Swartz, 2008*). To date, three-dimensional hERG structures have been determined by cryo-electron microscopy (cryo-EM), which has revealed that the VSD adopts the S4-up conformation under the nominal absence of membrane potential in detergent micelles (*Wang and MacKinnon, 2017*). By contrast, the resting-state structure, in which the VSD adopts the S4-down conformation, has not been determined because structural analysis under resting membrane potential is technically challenging.

Specific ligands that stabilize the resting state of hERG include APETx1, which is a 42-amino-acid peptide toxin of the sea anemone *Anthopleura elegantissima* (*Diochot et al., 2003*). APETx1 is a gating-modifier toxin that binds to the VSD and inhibits the voltage-dependent activation of hERG, thus stabilizing the resting-state, S4-down conformation of the VSD (*Diochot et al., 2003; Zhang et al., 2007*). Therefore, APETx1 is an effective tool for characterizing the molecular surface of the hERG VSD in the S4-down conformation, and for exploring the binding sites of the specific ligands that control hERG function.

A previous study, which conducted cysteine-scanning analysis of hERG, identified two residues in the S3-S4 region of the VSD that play important roles in hERG inhibition by APETx1 (*Zhang et al., 2007*). However, mutational analysis of APETx1 could not be conducted because only natural resources have been available until now; thus, no information could be obtained regarding the APETx1 residues crucial for hERG inhibition. Therefore, it remains unclear how APETx1 binds to the S4-down conformation of the VSD to inhibit hERG activation.

In the present study, we established a method of preparing recombinant APETx1 and investigated the hERG inhibition activity of APETx1 through electrophysiological analyses of APETx1 and hERG mutants. We identified the hydrophobic residues of APETx1 and in the hERG S3-S4 region that are related to hERG inhibition by APETx1. Our docking model well explains how APETx1 recognizes and stabilizes the S4-down conformation of the hERG VSD, and thus provides a structural basis for designing ligands that control hERG function by binding to the VSD.

## Results

### Functional and structural characterization of recombinant APETx1

In a previous study, APETx1 was purified from the sea anemone *Anthopleura elegantissima* (hereafter referred to as “natural product”) in order to characterize its inhibitory effect on hERG and its solution structure (*Chagot et al., 2005; Diochot et al., 2003; Zhang et al., 2007*). Here, we prepared recombinant APETx1, which was expressed as inclusion bodies in *E. coli*, purified in urea buffer, and refolded by dialysis. This dialysis process allowed the formation of three intramolecular disulfide bonds (see Materials and Methods for details of the sample preparation). We examined whether the prepared recombinant APETx1 is equivalent to the APETx1 isolated from the natural product in terms of structure and function.

First, a set of amide proton chemical shifts of recombinant APETx1 was compared with the corresponding set from the natural product (established at pH 3.0 and 280 K; *Chagot et al., 2005*) in order to examine their structural equivalence. We established the NMR resonance assignments of the recombinant APETx1 at pH 6.0 and 298 K (Supplementary Figure 1a-c), which were transferred to those at pH 3.0 and 280 K (Supplementary Figure 2a-d). It should be noted that the backbone amide signals of Y5, F33, and L34 were not observed in the ^1^H-^15^N HSQC spectrum at pH 3.0 and 280 K. The chemical shift values of backbone amide protons uniformly deviated by 0.12 ppm on average from those previously reported (Supplementary Figure 2e) (*Chagot et al., 2005*). These differences are probably due to systematic errors, such as from chemical shift referencing. Taking into consideration these systematic differences, the chemical shift differences are within ± 0.05 ppm, indicating that the structure of recombinant APETx1 is identical to that of the natural product.

Next, the hERG inhibition effect of the recombinant APETx1 was evaluated and compared with that of the natural product (*Diochot et al., 2003; Zhang et al., 2007*). We observed hERG K^+^ currents in the presence or absence of prepared APETx1 by whole-cell patch-clamp recordings and two-electrode voltage clamp (TEVC) recordings. The data obtained from whole-cell patch-clamp recordings (Figure 1a-d) demonstrated that the recombinant APETx1 effectively inhibited the hERG currents and shifted the half-maximal activation voltage (*V*_1/2_) values toward positive voltage in a dose-dependent manner. These results are consistent with those of previous studies (*Diochot et al., 2003; Zhang et al., 2007*). Similar results were obtained from TEVC recordings (Supplementary Figure 3a and 3c-d). The apparent dissociation constant (K_d_) values for the binding of APETx1 to hERG were estimated based on the data obtained from TEVC recordings, according to a previously reported method (*Zhang et al., 2007*). The fraction of uninhibited currents (*I*_Toxin_ / *I*_Control_), which was derived from the tail currents at weak depolarizing pulses (−30 mV) was reduced by incremental increases of APETx1, and the data were fitted with three models, models (A)-(C), as described previously (Supplementary Figure 3b) (*Zhang et al., 2007*). Because residual uninhibited currents were observed even at the 10 μM concentration of APETx1, models (A) and (C), which assume fractional toxin-sensitive currents, fitted the data more closely than model (B), which assumes a fully toxin-sensitive current; this is consistent with the findings of a previous study (*Zhang et al., 2007*). It is presently unclear which model supports APETx1 binding to hERG. It should be noted that the calculated *K*_d_ values of the three models are as follows: (A) 1.2 μM, (B) 1.7 μM, and (C) 0.23 μM; these are 12- to 14-fold higher than *K*_d_ values from a previous study, which are: (A) 87 nM, (B) 141 nM, and (C) 16.3 nM (*Zhang et al., 2007*). Differences between the *K*_d_ values in the previous and present studies are likely due to technical issues (see Discussion).

**Figure 1.**
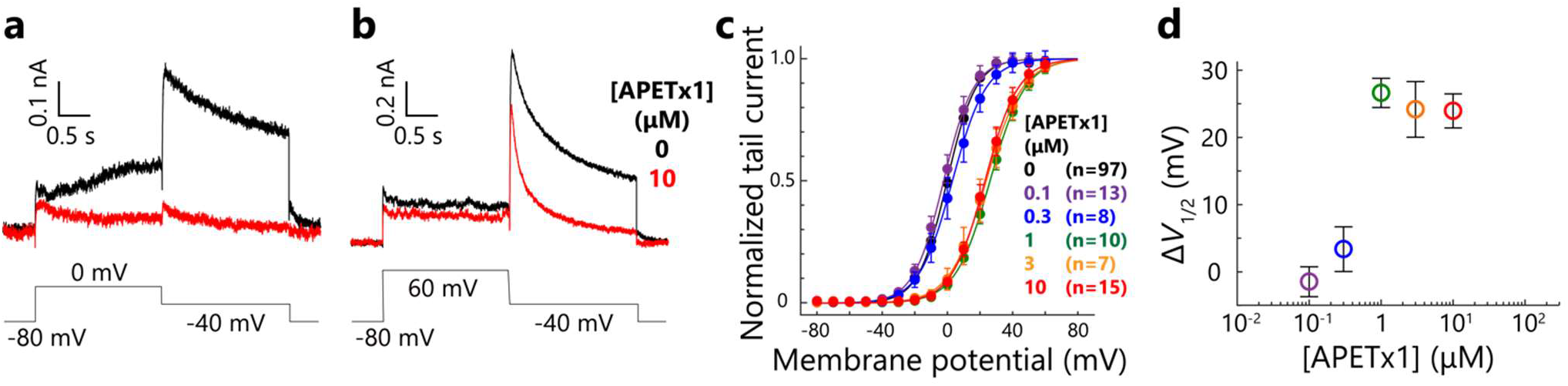
Functional and structural characterization of recombinant APETx1 in patch-clamp recordings. (a, b) Current traces of hERG elicited by two different depolarization pulses, 0 mV (a) and 60 mV (b), before (black) and after (red) the administration of 10 μM APETx1. Voltage protocol is illustrated at the bottom of each current trace. (c) Normalized *G*-*V* curves (mean ± SEM) of hERG in the presence or absence of different APETx1 concentrations. (d) The Δ*V*_1/2_ values of different concentrations of APETx1; these values indicate the toxin-induced shift of the half-maximal activation voltage of hERG.

Based on the NMR and electrophysiological results, we concluded that the recombinant APETx1 prepared in the present study displays properties (i.e., structure and hERG inhibitory activity) that are identical to those of the natural product used in previous studies (*Chagot et al., 2005; Diochot et al., 2003; Zhang et al., 2007*).

### Solution structure of APETx1 at pH 6.0

We observed pH-dependent chemical shift changes in the backbone-amide signals between pH 3.0 and pH 6.0 for the C-terminal residues V41 and D42 and the side-chain signal of R24 (Supplementary Figure 2f). These results suggest that the previously reported structure, determined at pH 3.0 (*Chagot et al., 2005*), might be different from that which occurs under physiological pH conditions. We therefore determined the three-dimensional structure of the recombinant APETx1 at pH 6.0 (Figure 2a and Table 1), and compared this with the structure at pH 3.0 (Figure 2b; *Chagot et al., 2005*). Backbone overlay of these two structures clearly shows that the structure of the C-terminal region is different (Figure 2b). At pH 6.0, the side-chain carboxyl group of D42 lies closer to the guanidinium group of R24, possibly because the deprotonation of the former enables the formation of hydrogen bonds or electrostatic interaction with the latter. pH-titration experiments showed that the changes in chemical shift values are reversible (data not shown). These results indicate that although the conformation of the C-terminal region is slightly altered according to pH conditions, the overall structure of APETx1 remains essentially identical.

**Figure 2.**
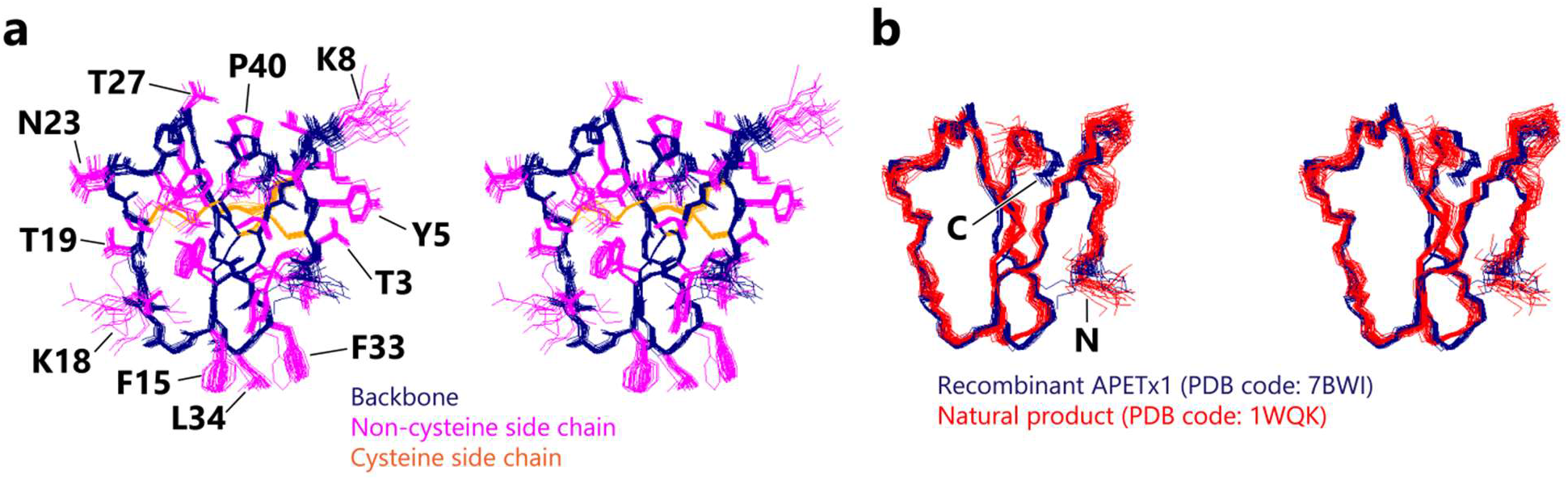
Structure determination of recombinant APETx1 at pH 6.0 and 298 K. (a) Stereo view of the ensemble of the 20 lowest-energy structures of recombinant APETx1, colored as follows: heavy atoms of backbone, navy; non-cysteine side chain, magenta; cysteine side chain, orange. (b) Stereo view of the overlay of the 20 structures of recombinant APETx1 at pH 6.0 and 298 K from the present study (PDB code: 7BWI; navy) and the 25 structures of the natural product at pH 3.0 and 280 K from a previous study (PDB code: 1WQK; red; *Chagot et al., 2005*).

**Table 1.** Structural statistics of the APETx1 structures in the present study (recombinant protein) and the previous study (natural product; *Chagot et al., 2005*).

|  | Natural product | Recombinant protein |
| --- | --- | --- |
| Experimental conditions | pH 3.0 and 280K | pH 6.0 and 298K |
| PDB code | 1WQK | 7BWI |
| Distance restraints |  |  |
| Total NOE-derived restraints | 751 | 766 |
| Intraresidue restraints ($\vert i-j\vert = 0$) | 366 | 131 |
| Sequential restraints ($\vert i-j\vert = 1$) | 140 | 216 |
| Short-range restraints ($2 \leq \vert i-j\vert \leq 4$) | 61 | 94 |
| Long-range restraints ($\vert i-j\vert \geq 5$) | 184 | 325 |
| Disulfide bond restraints | 9 | 12 |
| Dihedral angle restraints | 20 | 41 |
| Hydrogen-bond restraints | 36 | - |
| Root-mean-square deviation (RMSD) from mean coordinate structure (Å) |  |  |
| Backbone heavy atoms |  |  |
| Residues 1-42 | $0.82 \pm 0.17$ | $0.66 \pm 0.21$ |
| Residues 2-41 | $0.63 \pm 0.13$ | $0.48 \pm 0.12$ |
| All heavy atoms |  |  |
| Residues 1-42 | $1.28 \pm 0.17$ | $1.01 \pm 0.15$ |
| Residues 2-41 | $1.13 \pm 0.15$ | $0.95 \pm 0.14$ |

### Four clustered hydrophobic residues of APETx1 contributing to hERG inhibition identified by mutational analysis

To identify APETx1 residues that contribute to hERG inhibition, we selected 15 solvent-exposed residues (T3, Y5, K8, F15, K18, T19, S22, N23, R24, T27, S29, Y32, F33, L34, and D42) on the molecular surface of APETx1 for scanning mutational analysis (Figure 3a-b). We designed alanine-substitution mutants, omitting T3, as it has been reported that the natural T3P mutant, called APETx3, exhibits no hERG inhibition (*Peigneur et al., 2012*). This consideration motivated us to design a proline-substitution mutant. The ^1^H-^15^N HSQC spectra of all mutants are essentially identical to that of the wild-type (WT) APETx1; the exceptions to this are the mutated residue and a few neighboring residues (Supplementary Figure 4). This result indicates that none of the mutations affected the overall APETx1 structure.

**Figure 3.**
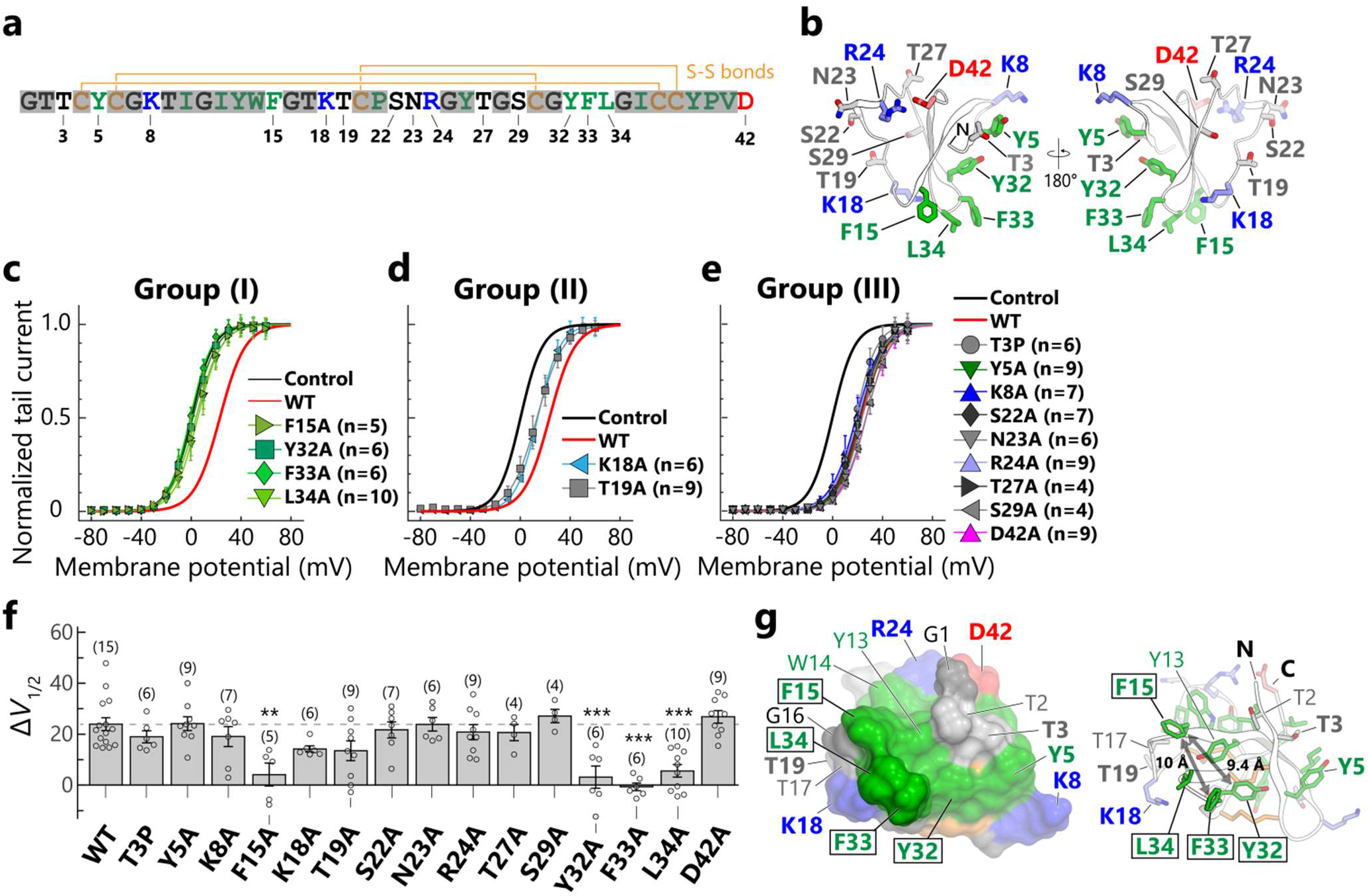
hERG inhibition by APETx1 mutants. (a) The primary structure of APETx1, colored as follows: positively-charged residues (K and R), blue; negatively-charged residues (D), red; hydrophobic residues (F, I, L, P, V, W, and Y), green; cysteine residues, orange; and others (G, N, S, and T), black. Three pairs of disulfide bonds are shown with orange lines. Non-mutated residues are drawn with semi-transparent gray marker to highlight the mutated residues. (b) The structure of APETx1 is shown as a ribbon representation with sticks depicting the mutated residues. (c-e) Normalized *G*-*V* curves (mean ± SEM) of hERG in the presence of 10 μM APETx1 mutants expressed by different colors and symbols. Control and WT correspond to the normalized *G*-*V* curve in the presence and absence, respectively, of 10 μM APETx1 from Figure 1c. Based on the Δ*V*_1/2_ values, APETx1 mutants were categorized into the groups (I)-(III). (f) The Δ*V*_1/2_ values of 10 μM APETx1 mutants in hERG are represented as the mean values ± SEM, with the number of experiments shown in parentheses. Multiple-group comparison was performed by one-way ANOVA followed by Tukey’s test (*, 0.01 ≤ *p* < 0.05; **, 0.001 ≤ *p* < 0.01; ***, *p* < 0.001). (g) Close-up view of the hydrophobic residues that yielded group (I) mutants (boxed residues) are shown as surface representations (left) and ribbon representations with sticks depicting side chains (right). The mutated residues are shown in bold. The distance between the Cβ atoms of F15 and Y32 is 9.4 Å, and that between F15 and F33 is 10 Å.

hERG currents were measured in the presence or absence of the APETx1 mutants by whole-cell patch clamp recordings (Figure 3c-f, Supplementary Figure 5). The toxin-induced *V*_1/2_ shift (Δ*V*_1/2_) value of 10 μM WT APETx1 was approximately 24 mV (Δ*V*_1/2_ = 23.9 ± 2.5 mV; Figure 1d). The APETx1 mutants were categorized into three groups according to the Δ*V*_1/2_ values: (I) mutants that showed significantly decreased Δ*V*_1/2_ values relative to those of WT (F15A, Y32A, F33A, and L34A; Figure 3c and 3f); (II) mutants exhibiting no significant change but a decreasing tendency in Δ*V*_1/2_ values relative to those of WT (K18A and T19A; Figure 3d and 3f); and (III) mutants showing Δ*V*_1/2_ values nearly equal to those of WT (T3P, Y5A, F15A, S22A, N23A, T27A, S29A, and D42A; Figure 3e-f). These results clearly indicate that F15, Y32, F33, and L34 play key roles in hERG inhibition. These four residues are localized on the molecular surface of APETx1, while K18 and T19, which yielded group (II) mutants, lie on the periphery of the group (I) site (Figure 3g). In contrast to these two groups, the residues that yielded group (III) mutants were not located close to the four key residues, but were instead dispersed on the molecular surface of APETx1 (Figure 3b). These results suggest that the molecular surface formed by the residues F15, Y32, F33, and L34 are foundational to the interactions between APETx1 and hERG.

As noted above, APETx3 is reported to show no inhibitory effect on hERG (*Peigneur et al., 2012*). Unexpectedly, our results showed that recombinant APETx3 shifted the *V*_1/2_ values toward positive voltage by about 19 mV, which is comparable to the effect of APETx1 (Figure 3f). It is worth noting that the *V*_1/2_-shift effect of the natural-product APETx3 has not been characterized, and that preparation methods and experimental conditions in the present study are different from those in the previous study (*Peigneur et al., 2012*).

### Mutations of hERG residues that affect inhibitory activity of APETx1

We introduced a mutation to hERG to investigate whether this would affect the inhibitory activity of APETx1 and thereby identify the APETx1-binding site on hERG. We proceeded based on the generally accepted proposal that the extracellular side of the VSD of a voltage-gated ion channel is targeted by gating-modifier toxins (*Swartz, 2007*). A previous study examining the APETx1-hERG interaction via cysteine-scanning mutational analysis of G514-E519, which are located in the S3-S4 region of the VSD, showed that G514C and E518C mutations respectively increased and decreased the Δ*V*_1/2_ value of 10 μM APETx1 (*Zhang et al., 2007*). These results suggest that these hERG residues in the S3-S4 region are involved in APETx1 binding (*Zhang et al., 2007*). As mentioned above, APETx1 shifted the *V*_1/2_ values toward positive voltage (Figure 1c-d and Supplementary Figure 3c-d), which supports the possibility that APETx1 preferentially binds to hERG in its resting state, as described previously (*Diochot et al., 2003; Zhang et al., 2007*). However, it remains unknown where and how APETx1 interacts with the VSD to stabilize the resting state, S4-down conformation of hERG. Therefore, in the present study, we conducted a mutational analysis of the residues in the S3-S4 region of hERG, focusing on F508-L524, and omitting S515-S517 and E519, which have been previously investigated (Figure 4a-c; *Zhang et al., 2007*). We also examined the mutational effect of L433 and the charged residues (K434, E435, E437, E438, D456, and D460) on the S1-S2 region to confirm whether the S1-S2 region contributes to hERG inhibition of APETx1 (Figure 4a-b and 4d).

**Figure 4.**
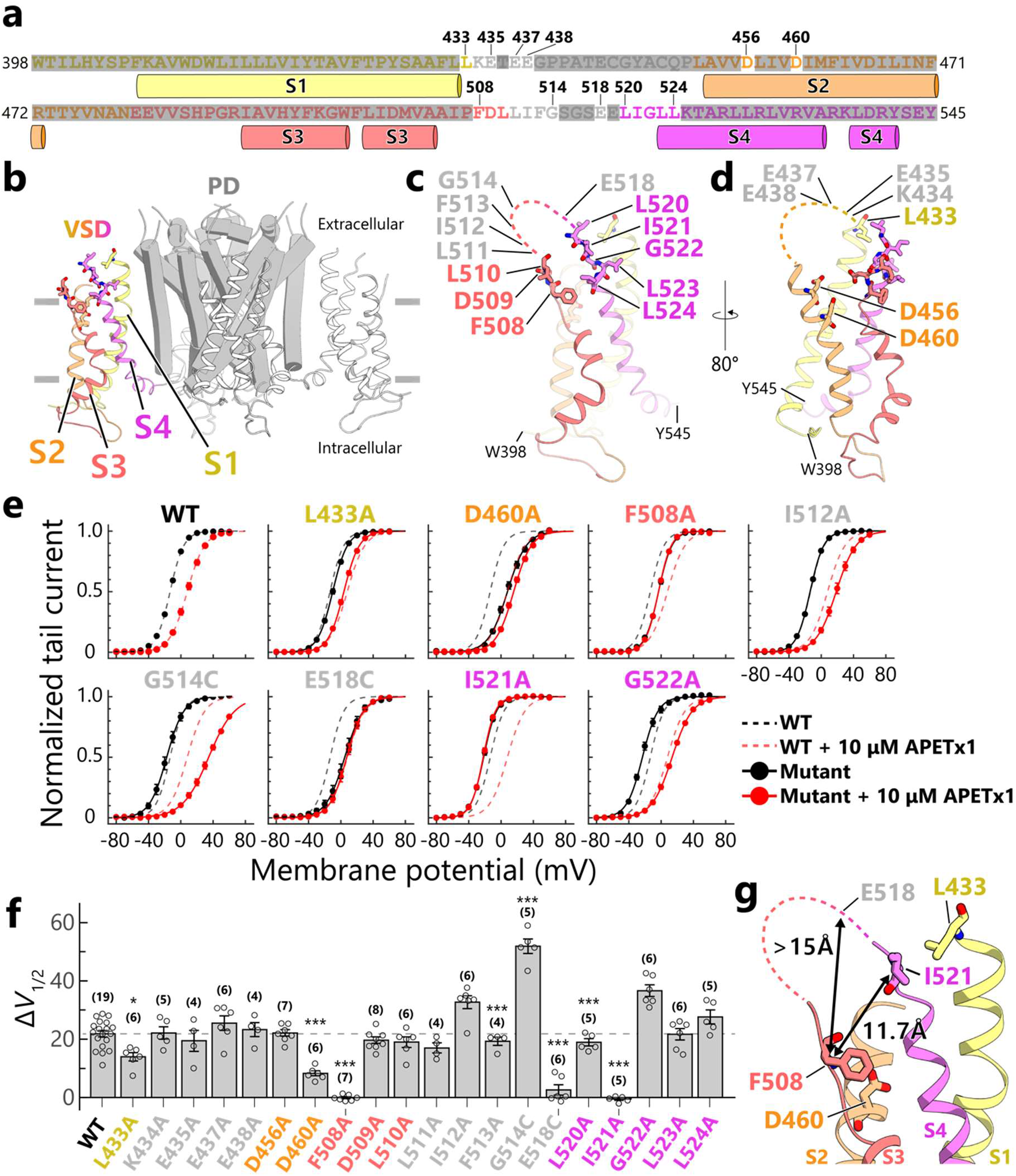
Inhibition of hERG mutants by APETx1. (a) The primary structure of the hERG VSD, colored as follows: S1, yellow; S2, orange; S3, pale red; and S4, magenta. Cylinders attached below the sequence indicate helical segments. The residues that are missing in a cryo-EM structure (PDB code 5VA2) are colored gray. Non-mutated residues are drawn with semi-transparent gray marker to highlight the mutated residues. (b) The transmembrane domain of the cryo-EM structure of hERG with S4-up conformation (PDB code: 5VA2; *Wang and MacKinnon, 2017*). The VSDs are represented as ribbons, one of which is colored as in (a); the others are white. The PD is represented as a gray cartoon. (c, d) A detailed illustration of the hERG VSD, corresponding to the sequence shown in (a). The mutated residues are represented as sticks. The residues in the extracellular region that are missing in the cryo-EM structure of hERG (PDB code: 5VA2) are depicted as dashed lines. (e) Normalized *G*-*V* curves (mean ± SEM) of hERG mutants showing Δ*V*_1/2_ values that are significantly different from those of WT, in the absence (black circle and solid line) or presence (red circle and solid line) of 10 μM APETx1. The fitting curves of WT in the presence (black dashed line) and absence (red dashed line) of 10 μM APETx1 are superimposed on those of the mutants. (f) The Δ*V*_1/2_ values of 10 μM APETx1 in hERG-mutants are represented as the mean values ± SEM; the number of experiments is shown in parentheses. Multiple-group comparison was performed by one-way ANOVA followed by Tukey’s test (*, 0.01 ≤ *p* < 0.05; **, 0.001 ≤ *p* < 0.01; ***, *p* < 0.001). (g) Close-up view of the residues that yielded mutations that decreased the hERG inhibition by APETx1. The distance between the Cβ atoms of F508 and I521 (indicated by an arrow) are 11.7 Å, and the distance between F508 and E518 is at least 15 Å.

We measured the currents of the hERG mutants by TEVC recordings using oocytes from *Xenopus laevis*. All mutants expressed in *X. laevis* oocytes showed hooked tail currents, which are characteristic features of hERG (Supplementary Figure 6-9; *Vandenberg et al., 2012*). This demonstrates that the overall structures of hERG mutants are not substantially different from those of WT. It should be noted that, even in the absence of APETx1, some mutants (D456A, D460A, D509A, E518C, and L524A) exhibited *V*_1/2_ shifts toward positive voltage and others (L520A and L523A) toward negative voltage (Supplementary Figure 6-9). Therefore, the results pertaining to these mutants should be carefully evaluated.

We found that the Δ*V*_1/2_ values of 10 μM APETx1 in TEVC recordings were comparable to those in patch-clamp recordings in WT hERG (Δ*V*_1/2_ = 21.9 ± 1.1 mV, in TEVC recordings; Δ*V*_1/2_ = 23.9 ± 2.5 mV, in patch-clamp recordings; Figures 1d and 4e). We confirmed that the Δ*V*_1/2_ values were increased by the G514C mutation and decreased by the E518C mutation (Figure 4f), which is consistent with the results of a previous study (*Zhang et al., 2007*). We further found that the F508A and I521A mutations significantly decreased the Δ*V*_1/2_ values relative to WT, while the I512A and G522A mutations increased the values (Figure 4c and 4e-f). Furthermore, mutations of L433 on S1 and D460 on S2 also significantly decreased the Δ*V*_1/2_ values relative to WT (Figure 4d-f). These results suggest that the S3-S4 loop plays a key role in APETx1 binding, while L433 on S1 and D460 on S2 are also involved in the hERG inhibition of APETx1.

## Discussion

### Structure and function of the recombinant APETx1

In the present study, we conducted structural and electrophysiological analyses of recombinant APETx1 and its mutants. We concluded that it is structurally identical to the natural product based on the chemical shift values of each under identical conditions (pH 3.0, 280 K; Supplementary Figure 2e). The structure of the C-terminal region determined in the present study at pH 6.0 and 298 K was slightly different from that of the natural product determined in a previous study under different conditions (pH 3.0, 280 K; *Chagot et al., 2005*). We attribute this difference to variations in pH (Supplementary Figure 2f), and we used ^1^H-^15^N HSQC spectra to confirm that this structural change is reversible.

Recombinant APETx1 effectively inhibited the hERG currents elicited by relatively weak depolarization pulses (Figure 1a), shifting the *V*_1/2_ values toward positive voltage in a dose-dependent manner (Figure 1c-d, Supplementary Figure 3c-d). This positive *V*_1/2_-shift effect suggests that APETx1 recognizes and stabilizes the resting state of hERG, in which the VSD adopts the S4-down conformation, as described previously (*Diochot et al., 2003; Zhang et al., 2007*). It should be noted that APETx1 caused a reduction in the maximal conductance at higher voltages and faster attenuation of the tail currents (Figure 1b and Supplementary Figure 6), suggesting that APETx1 remains hERG-bound even when strong depolarizing pulses are applied. These implicative phenomena were also reported in several other gating-modifier toxins that stabilize the S4-down conformation of voltage-gated ion channels (*Ebbinghaus et al., 2004; Herrington, 2007; Lee et al., 2004; Li-Smerin and Swartz, 1998; Phillips et al., 2005; Shen et al., 2019; Swartz, 2007; Swartz and MacKinnon, 1997; Xu et al., 2019*).

A previous study reported that an increase in Δ*V*_1/2_ values occurs at a micromolar concentration of APETx1 (10^−6^ M), while the *K*_d_ values derived from *I*_toxin_ / *I*_control_ at weak depolarizing pulses are 10^−7^-10^−8^ M (*Zhang et al., 2007*). In our TEVC recordings, however, both a decrease in *I*_toxin_ / *I*_control_ (Supplementary Figure 3b) and an increase in the Δ*V*_1/2_ values (Supplementary Figure 3d) were observed at micromolar concentrations. The reasons for the differences between the present and previous studies remain unclear; however, they might be partially due to the following technical issues that made accurate determination of the *K*_d_ values difficult (*Zhang et al., 2007*): (1) inaccuracy of small current amplitudes at weak threshold depolarization pulses, and (2) variations among oocytes in responsiveness to APETx1. Therefore, we evaluated the Δ*V*_1/2_ values in the presence of APETx1 at a single concentration (10 μM) for the mutational analyses of APETx1 and hERG, according to a previous study (*Zhang et al., 2007*).

### Hydrophobic surface of APETx1 contributes to hERG inhibition

We established a method for the preparation of recombinant APETx1, which enabled us to conduct mutational analysis. The electrophysiological analysis using the APETx1 mutants clearly showed that the four hydrophobic residues (F15, Y32, F33, and L34) play pivotal roles in hERG inhibition. Two hydrophilic residues, K18 and T19, also appear to contribute to the inhibition. By mapping these residues onto the APETx1 structure, we revealed that they are localized on the APETx1 molecular surface (Figure 3g), where the four hydrophobic residues are clustered at the edge of the large hydrophobic surface; this is shown in green in Figure 3g. It is well known that many gating-modifier toxins possess a large hydrophobic surface, called a hydrophobic patch, which is reported to play a role in partitioning the cell membrane prior to the binding of the target channel (*Agwa et al., 2017*). The hydrophobic patch on APETx1 also appears to contribute to such membrane-partitioning, although this has not been reported for APETx1. The mutation of F15, Y32, F33, or L34 to alanine, which possesses a small but hydrophobic side chain, probably has only a limited effect on the membrane-partitioning capacity of APETx1. Therefore, the drastic decreases observed in the Δ*V*_1/2_ values of the mutations of the four residues, along with their localization on the APETx1 surface, strongly suggest that these residues engage in direct interactions with hERG in the membrane, which could stabilize the hERG resting state.

### Putative APETx1-binding residues of hERG mapped onto the S4-down conformation

In the present study, we identified four previously undescribed hERG residues in the S3-S4 region (F508, I512, I521, and G522); mutations of these, along with the previously reported residues G514 and E518 (*Zhang et al., 2007*), affect hERG inhibition of APETx1 (Figure 4f). These results show that the structure of the S3-S4 region plays a key role in APETx1-binding.

One of the best ways to obtain pairwise information on the interacting residues between hERG and APETx1 is through thermodynamic mutant cycle analysis (*Hidalgo and MacKinnon, 1995*). However, this analysis could not be conducted in the context of the hERG-APETx1 interaction because even a single mutation in either APETx1 or hERG decreases the Δ*V*_1/2_ values to nearly zero in the presence of 10 μM APETx1 or its mutant (Figure 3f and 4f); thus, the additive effect of the double mutation cannot be evaluated. Therefore, we speculate on an APETx1-hERG interaction mode, as described below.

The Cβ atoms of the four hydrophobic residues (F15, Y32, F33, and L34) of APETx1 are all located within 10 Å of each other (Figure 3g), suggesting that the interacting counterpart residues are complementarily distributed on the hERG molecular surface. In the hERG S4-up conformation, which was revealed by cryo-EM analysis (*Wang and MacKinnon, 2017*), the distance between the Cβ atoms of F508 and I521 is approximately 12 Å, and that between F508 and E518 appears to be greater than 15 Å (Figure 4g); such distances are not complementary to the distribution of the APETx1 active residues. We then built structural models of the S4-down conformation of the hERG VSD and investigated whether the models are consistent with our mutational analysis results.

Among the residues, E518 is essential for the inhibition of APETx1, pointing to the possibility that the side-chain carboxyl group is important for direct interaction with APETx1 (Figure 4f; *Zhang et al., 2007*). We mutated the following APETx1 residues, which are exposed on its molecular surface, as candidates for residues that interact with hERG E518: the basic residues (K8, K18, and R24) and those with hydrogen-bonding ability (T3, Y5, T19, S22, N23, T27, S29, and Y32). Of these, only the Y32A mutation significantly decreased hERG inhibition by APETx1 (Figure 3f). Therefore, we speculate that Y32 of APETx1 directly interacts with E518 of hERG. The fact that E519, the residue next to E518, is not involved in the hERG inhibition of APETx1 (*Zhang et al., 2007*) indicates that APETx1 distinguishes between E518 and E519. Although these two acidic residues are missing in the cryo-EM structure of hERG (*Wang and MacKinnon, 2017*), we assumed that they adopt a particular conformation, with their carboxyl groups pointing in different directions. Indeed, the cryo-EM structure of the closely related rat KV channel, rEAG1 (*Whicher and MacKinnon, 2016*), reveals that the position corresponding to the hERG E518-E519 residues adopts a helical structure that extends from an S4 helix (Supplementary Figure 10a).

In order to visualize the binding mode of APETx1 to hERG, we used the structural model of rEAG1 as a template (*Whicher and MacKinnon, 2016*) for the construction of an S4-up model (“up” model, Supplementary Figure 10b). We further constructed two hypothetical S4-down models, the “one-helical-turn down” model and the “two-helical-turn down” model (Supplementary Figure 10b), in which the S4 helix is shifted toward the intracellular side by one and two helical turns, respectively. These S4-down models are based on the premise that in response to membrane repolarization, the S4 helix moves toward the intracellular side, with the basic residues on the S4 helix maintaining salt bridges with the acidic residues on the S1-S3 helices (*Bezanilla, 2008; Börjesson and Elinder, 2008; Groome and Bayless-Edwards, 2020; Swartz, 2008; Vargas et al., 2011*).

In these models, the side chains of hERG E518 and I521 are exposed and constitute the largest surface areas in the S4-down models (Supplementary Figure 10e and 10g), which is in contrast to the S4-up model (Supplementary Figure 10c). The S4-down models show that the key hERG residues (F508, E518 and I521) are localized and form a crevice on the molecular surfaces, which appears complementary to the distribution of the APETx1 active residues (Supplementary Figure 10f and 10h, left, white dotted circle). By contrast, the S4-up model shows that the key hERG residues are not localized and that no crevice is formed on the molecular surface, suggesting that all APETx1 active residues cannot simultaneously interact with the hERG S4-up conformation (Supplementary Figure 10d).

### Docking of APETx1 and the S4-down conformation of hERG-VSD

Next, we built the structural models of the APETx1-hERG complex to account for the results of the mutational analyses in the previous (*Zhang et al., 2007*) and present studies. In these models, APETx1 is docked on the S4-down models based on the assumptions that the key residues, identified by the mutational analyses of APETx1 and hERG, interact with each other directly, and that hydrogen bond form between the side chains of APETx1 Y32 and hERG E518 (Figure 5). It should be noted that S3 and S4 of the hERG VSD face the hydrophobic region of the lipid bilayer; thus, when bound to the hERG VSD, APETx1 is reasonably accommodated on S4, without contacting the PD or other VSDs (Supplementary Figure 11a-b). We also performed a docking simulation of APETx1 on the S4-up model. This resulted in a complex model with no surface complementarity between the identified APETx1 and hERG residues (data not shown). This suggests that the mutational effects on hERG inhibition by APETx1 predominantly reflect a change in the interaction between APETx1 and the S4-down conformation, rather than the S4-up conformation.

Figure 5 shows the mapping of the residues that are key to hERG inhibition by APETx1 (see also Supplementary Figure 10c-h). On the S4-down models, a crevice formed by F508, E518, and I521 accommodates the F15, F33, and L34 side chains of APETx1 (Figure 5a-b and Supplementary Figure 10f and 10h). Not only are the distributions of these residues on each molecule complementary in these models, but the shapes of the molecular surfaces are as well. However, we were unable to determine which of the two S4-down models is more reasonable.

**Figure 5.**
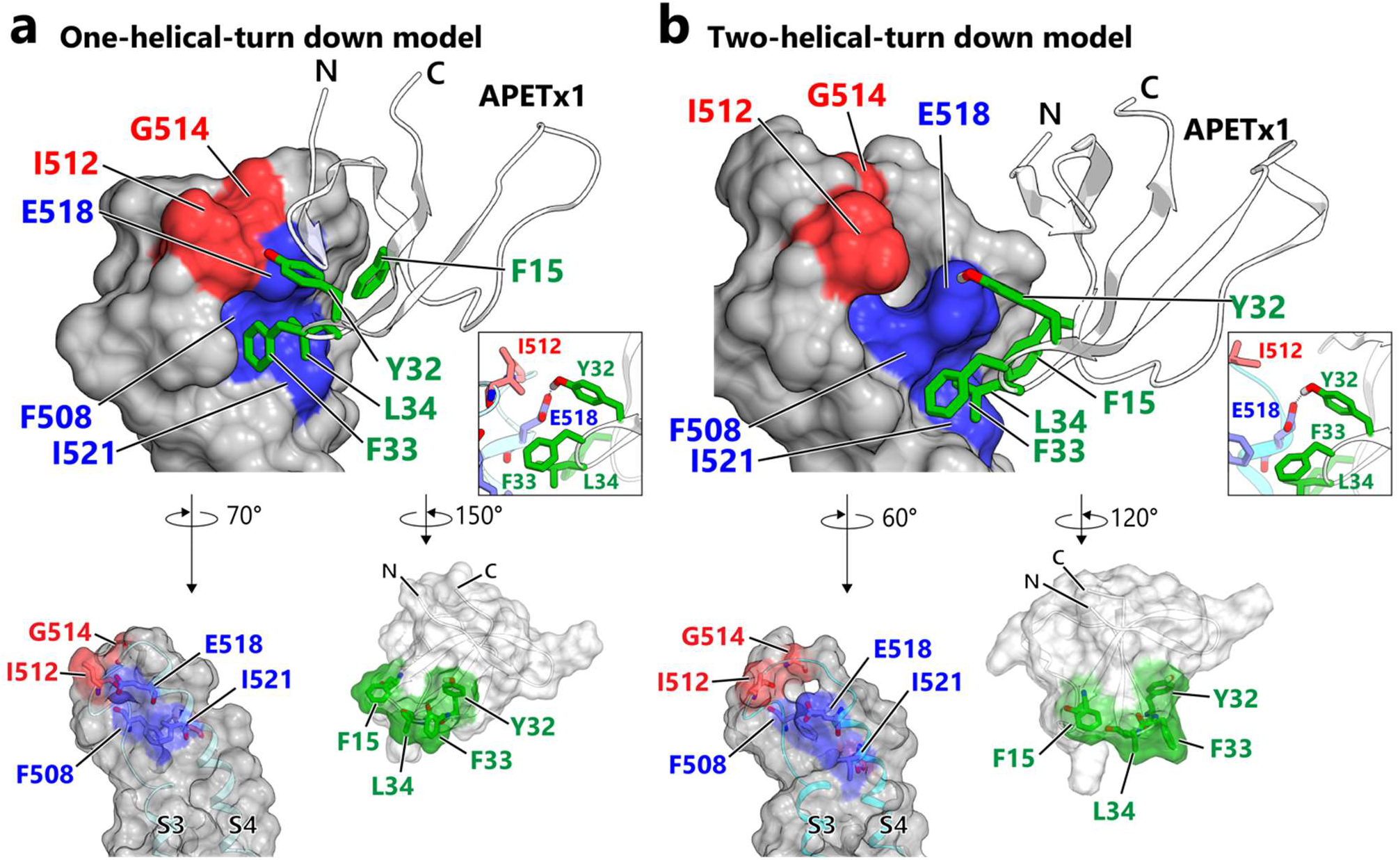
Structural models of the APETx1-VSD complex that satisfies the results of the mutational analysis. APETx1 docked to the one- and two-helical-turn-down model of hERG VSD in (a) and (b), respectively. The models exclude the S1 and S2 regions, which do not come into contact with APETx1, for clarity. The models of the hERG S3-S4 regions are represented as molecular surfaces colored according to the residues that yielded mutations that decreased (blue) and increased (red) inhibition by APETx1. APETx1 is represented as a ribbon with green sticks representing the key residues involved in hERG inhibition. Close-up view of the hydrogen bond between APETx1 Y32 and hERG E518 is shown in inset. “Open-book” representations of the interaction interfaces are also drawn as semi-transparent molecular surfaces with ribbon representation.

APETx1 showed stronger inhibition of the hERG I512A mutant than of the WT (Figure 4f). In the one-helical-turn down model, the sterically bulky I512 side chain lies in close proximity to E518 (Figure 5a, inset), suggesting that I512 hinders the access of APETx1 to E518 in the S4-down conformation. The increased inhibition of the I512A mutant by APETx1 could be due to the removal of the steric hindrance, resulting in a higher binding affinity to the S4-down conformation. Moreover, the mutants of the glycine residues of hERG, G514C (*Zhang et al., 2007*) and G522A, exhibit increased inhibition by APETx1 (Figure 4f). These remarkable mutational effects might be due to the structural changes caused by the mutation of the glycine residues adjacent to the important residues (e.g., I512 and E518), which optimizes the contact surface for APETx1 binding.

In the present study, recombinant APETx3 showed a *V*_1/2_ value comparable to that of APETx1 (Figure 3f). Although it is uncertain whether this result can be applied to the natural-product APETx3, the T3 of APETx1 has little involvement in hERG inhibition; this is consistent with our model, in which T3 of APETx1 does not come into direct contact with the hERG VSD.

The mutational analysis of hERG clearly shows that the S3-S4 region is crucial for inhibition by APETx1, but we cannot rule out the possibility that an additional hERG region is also involved in binding with APETx1. In particular, the mutations of L433 on S1 and D460 on S2 decreased inhibition by APETx1 (Figure 4f-g). Although these residues do not make direct contact with APETx1 in our models (Supplementary Figure 11a-b), their mutations might affect the structure of the APETx1-binding site in the S3-S4 region.

## Conclusion

The present study identified the key residues of APETx1 and hERG that are involved in hERG inhibition by APETx1. We built structural models that satisfy the electrophysiological results, in which APETx1 binds to the S4-down conformation of the hERG VSD, based on the assumption that the distribution of these residues on each molecule is complementary. These models will help advance understanding of the inhibitory mechanism of APETx1 and other gating-modifier toxins, and provide a structural basis for the creation of novel types of drugs targeting the VSDs of K_V_ channels.

## Materials and Methods

### Recombinant expression, purification, and refolding of WT and mutated APETx1

The DNA sequence encoding APETx1, along with an upstream TEV-protease-recognition sequence, were cloned into the pET-30Xa/LIC vector (Novagen). All APETx1 mutants were generated by PCR-mediated site-directed mutagenesis and confirmed by DNA sequence analysis.

The plasmids were transformed into Escherichia coli strain C41 (λDE3) for recombinant protein production. The cells were grown in Luria-Bertani medium supplemented with 40 mg/mL kanamycin, and maintained at 37°C with shaking at 150 rpm to an optical density of 0.6-1.0 at 600 nm. Expression of the His_6_-tag-fusion proteins was induced with 1 mM isopropyl β-D-1-thiogalactopyranoside (IPTG); the cells were grown for 6-12 h at 37°C and then harvested by centrifugation for 10 min at 5000×g. Uniformly ^15^N-labelled or uniformly ^13^C- and ^15^N-labelled APETx1 samples were prepared for NMR experiments by growing *E. coli* cells in M9 minimal medium supplemented with ^15^NH_4_Cl or ^15^NH_4_Cl and ^13^C_6_ glucose.

The cells were harvested by centrifugation and disrupted by sonication in lysis buffer (20 mM Tris-HCl, pH 8.0, and 200 mM NaCl) supplemented with 0.5 mM 4-(2-aminoethyl) benzenesulfonyl fluoride hydrochloride, 0.15 μM aprotinin, 1 μM E-64, and 1 μM leupeptin. The cell pellets were prepared by centrifugation for 30 min at 10000×g. The pellets were resuspended in lysis buffer containing 0.1% NP-40, followed by centrifugation; this procedure was repeated three times for the removal of nucleic acid. The pellets were solubilized using the lysis buffer containing 8.0 M urea for 1-2 h at room temperature (25°C-27°C). After centrifugation for 30 min at 10000×g, the His_6_-tag-fusion proteins were purified from the supernatant by a HIS-Select^®^ Nickel Affinity Gel (Sigma) column. The eluted His_6_-tag-fusion proteins were reduced by 2 mM dithiothreitol (DTT), followed by dilution to 3 μM or less so as to avert aggregation during the subsequent refolding step. The reduced proteins were refolded by dialysis against a redox buffer (3 mM reduced glutathione, 0.3 mM oxidized glutathione, 10% glycerol, 20 mM Tris-HCl, pH 9.0, and 200 mM NaCl) until the concentration of urea was less than 200 mM. This internal solution was next dialyzed against a refolding buffer (20 mM Tris-HCl, pH 9.0, and 200 mM NaCl) until the urea concentration was less than 10 mM. The internal solution was concentrated using Sep-Pak^®^ C18 cartridges (Waters), and then lyophilized. The sample was dissolved in lysis buffer and then centrifuged to remove the aggregated proteins, which form intermolecular disulfide bonds. After digestion with His-tag-fused TEV protease at 25°C, cleaved His-tag and His-tag-fused TEV protease were removed using a HIS-Select^®^ Nickel Affinity Gel (Sigma-Aldrich) column. APETx1 was further purified by reverse-phase high-performance liquid chromatography (RP-HPLC) using an ODS-AM column (YMC). The eluted APETx1 was detected by matrix-assisted laser-desorption ionization–time-of-flight mass spectroscopy (MALDI-TOF MS) because APETx1 was only lightly stained in SDS-PAGE gel with coomassie brilliant blue (CBB).

### NMR resonance assignments of APETx1 and validation of mutants

Data were collected using Bruker Avance 500 or 600 spectrometers equipped with triple-resonance probes. For NMR resonance assignments of APETx1, we measured ^1^H-^15^N HSQC spectra, ^1^H-^13^C HSQC spectra, and non-uniformly sampled, three-dimensional NMR spectra using 258 μM uniformly ^13^C- and ^15^N-labelled APETx1 in a buffer containing 20 mM KH_2_PO_4_ (pH 6.0), 100 mM NaCl, and 10% D_2_O (hereafter referred to as “phosphate buffer”) at 298 K. The mixing times of the ^15^N-edited and the ^13^C-edited NOESY experiments, which were used for sequential assignments, were set to 100 ms and 120 ms, respectively. ^1^H-^15^N HSQC spectra were also measured at different pH levels (pH 3.0, pH 4.5, and pH 6.0; the phosphate buffer was pH-adjusted with HCl) and temperatures (280 K, 290 K, and 298 K) using 140 μM uniformly ^15^N-labelled APETx1.

The ^1^H-^15^N HSQC spectra of uniformly ^15^N-labelled WT APETx1 and selected mutants (T3P, K8A, F15A, K18A, R24A, Y32A, and F33A) were measured in the phosphate buffer at 298 K; those of WT and the remaining mutants (Y5A, T19A, S22A, N23A, T27A, S29A, L34A, and D42A) were measured in 10% D_2_O (pH 6.0) at 298 K. Sample concentrations of WT and mutants are as follows: WT, 149 μM (in the phosphate buffer) or 102 μM (in 10% D_2_O, pH 6.0); T3P, 50 μM; Y5A, 229 μM; K8A, 30 μM; F15A, 184 μM; K18A, 31 μM; T19A, 290 μM; S22A, 102 μM; N23A, 76 μM; R24A, 19 μM; T27A, 129 μM; S29A, 86 μM; Y32A, 408 μM; F33A, 634 μM; L34A, 258 μM; and D42A, 174 μM.

All spectra were processed using Bruker TopSpin 3.6 software or NMRPipe (*Delaglio et al., 1995*), and the data were analyzed with Sparky (T. D. Goddard and D. G. Kneller, Sparky 3, University of California, San Francisco, CA). The APETx1 backbone and side-chain NMR signals were sequentially assigned using non-uniformly sampled data for the following experiments: HNCACB, CBCA(CO)NH, HNCO, HCCH-COSY, HCCH-TOCSY, ^15^N-edited TOCSY, ^15^N-edited NOESY, and ^13^C-edited NOESY experiments. The amide signals of T2 and L34 were not sequentially assigned due to the absence of these signals on three-dimensional triple-resonance NMR spectra at pH 6.0; however, they were observed on ^1^H-^15^N HSQC spectra under low pH conditions (pH 3.0 and pH 4.5 at 298 K). ^1^H chemical shift values of WT APETx1 were obtained using sodium 4,4-dimethyl-4-silapentane-1-sulfonate (DSS) as a standard; ^13^C and ^15^N chemical shift values were also corrected indirectly. The ^1^H, ^13^C, and ^15^N chemical shift assignments at pH 6.0 and 298 K have been deposited in Biological Magnetic Resonance Bank (BMRB ID: 36345).

### NMR structure calculation of APETx1

Data were collected using Bruker Avance III HD 700 spectrometers equipped with triple-resonance cryogenic probes at 298 K. All experiments were performed using 691 μM uniformly ^13^C/^15^N-labelled APETx1 in a buffer solution containing 20 mM KH_2_PO_4_ (pH 6.0), 100 mM NaCl, and 10% D_2_O. The mixing times for the ^15^N-edited and the ^13^C-edited NOESY experiments for structural determination were set to 200 ms. These spectra were processed using NMRPipe (*Delaglio et al., 1995*).

Based on the chemical shift difference between Cβ and Cγ (*Schubert et al., 2002*), we confirmed that the peptide bond of the P40 residue is cis-conformer, which is consistent with the structure of the natural product (*Chagot et al., 2005*). This cis-peptide restraint was used for structure calculation. The dihedral angle restraints were predicted using TALOS+ and were based on the chemical shifts of ^13^Cα, ^13^Cβ, ^13^CO, ^15^Nα, and ^1^HN (*Shen et al., 2009*). NOE peaks were automatically picked using MagRO-NMRView (*Kobayashi et al., 2012; Kobayashi et al., 2018; Kobayashi et al., 2007*). The NOE peak intensity was converted to distance constraints, and structure calculation was performed using the torsion angle dynamics program CYANA 3.98 (*Guntert, 2004*). First, we calculated the preliminary structures without disulfide bond restraints to confirm that three pairs of cysteine residues can be correctly formed into disulfide bonds. Next, we calculated the structures using disulfide bond restraints and obtained 100 structures in the final iteration. The 20 structures with the lowest target function were refined by restrained molecular dynamics of 30 ps with Amber 12 (*Case et al., 2012*). The automated identification and superposition of the ordered regions of the determined structures were performed by using FitRobot ver. 1.00.07 (*Kobayashi, 2014*). Atomic coordinates of APETx1 and all restraint files used for the structure calculations have been deposited in the Protein Data Bank (PDB code: 7BWI). NMR structure ensembles were visualized and the root mean squared deviation (RMSD) values were calculated using MOLMOL (*Koradi et al., 1996*).

### Cell preparation for patch-clamp recordings

Stable HEK 293 cell lines expressing hERG (SB-HEK-hERG; SB Drug Discovery Limited) were used. The cells were cultured in Dulbecco’s Modified Eagle’s Medium (DMEM; Thermo, Gibco) supplemented with 10% fetal bovine serum (FBS; Thermo, Gibco) and 1% penicillin/stereptomycin (Thermo, Gibco) in a humidified 5% CO_2_ atmosphere at 37°C. For patch-clamp experiments, cultured cells on a polystyrene culture dish (Sumitomo Bakelite, Tokyo, Japan) were detached by TrypLE Express (Thermo, Gibco) at 37°C.

### Automated patch-clamp recordings for mutational analysis of APETx1

Whole-cell automated patch-clamp recordings were obtained using a SyncroPatch 384PE (Nanion Technologies GmbH, Germany) with single-hole medium resistance (4-5.5 MΩ) borosilicate glass planar chips. Pulse generation and data collection were performed with PatchControl 384 V1.6.6 and DataControl384 V1.8.0 software. Currents were sampled at 1 kHz. Leak subtraction was performed based on a small voltage step at the beginning of the voltage protocol. Seal resistance was calculated using built-in protocols, and cells with a seal resistance of 0.3-3 GΩ were analyzed.

For automated patch-clamp recordings, the intracellular solution contained 110 mM KF, 10 mM NaCl, 10 mM KCl, 10 mM EGTA, and 10 mM HEPES-KOH (pH 7.2, 285 mOsm); the extracellular solution contained 140 mM NaCl, 4 mM KCl, 2 mM CaCl_2_, 1 mM MgCl_2_, 5 mM D-glucose, and 10 mM HEPES-NaOH (pH 7.4, 298 mOsm). The command voltage step took into account the fact that the use of these solutions results in ~9 mV liquid junction potential. All experiments were performed at room temperature (20°C-25°C). The holding membrane potential was set at −80 mV. For the conductance-voltage (*G*-*V*) relationship experiments, hERG currents were elicited by depolarizing voltage steps from −80 mV to +60 mV in 10 mV increments for 2 s, followed by a step pulse to −40 mV for 2 s.

In order to determine the dose-response relationship, APETx1 was added to the extracellular solution at concentrations of 0.1-10 μM. APETx1 mutants were added to the extracellular solution at 10 μM for screening. The voltage-pulse protocol for determining the *G*-*V* relationship was performed after the effects of APETx1 WT or mutants reached a steady state.

### Ethical approval

All animal experiments were approved by the Animal Care Committee of the National Institutes of Natural Sciences (an umbrella institution of National Institute for Physiological Sciences, Tokyo, Japan), and were performed in accordance with its guidelines.

### Preparation for TEVC recordings

hERG was subcloned into an pSP64 plasmid, and hERG mutants were generated by PCR-mediated site-directed mutagenesis using an In-Fusion HD Cloning Kit (Takara, Otsu, Japan) as described previously (*Kume et al., 2018*). The DNA sequences of all mutants were confirmed by sequencing. The complementary RNAs (cRNA) were transcribed from each linearized plasmid DNA using the mMessage mMachine SP6 Transcription Kit (Ambion, Austin, TX, USA). *Xenopus laevis* were purchased from Hamamatsu Animal Supply Co. (Hamamatsu, Japan) and used for oocyte collection. Preparation and injection of X. laevis oocytes were performed as described previously (*Kume et al., 2018; Nakajo and Kubo, 2014*). Currents were measured 1-3 d after the cRNA injection, depending on the required current amplitude.

### TEVC recordings for mutational analysis of hERG

TEVC recordings were performed essentially as described previously (*Kume et al., 2018; Nakajo and Kubo, 2014*). Macroscopic currents were recorded from injected oocytes under a two-electrode voltage clamp (TEVC) using an amplifier (OC-725C; Warner Instruments, Hamden, CT, USA), an AD-DA converter (Digidata version 1440A; Molecular Devices, Sunnyvale, CA, USA), and software for control and recording of the voltage clamp (pCLAMP version 10.7; Molecular Devices). Glass microelectrodes were drawn from borosilicate glass capillaries (Harvard Apparatus, Cambridge, MA, USA) and filled with 3 M K-acetate and 10 mM KCl (pH 7.4 adjusted with HCl). The electrode resistance was 0.2–0.8 M. Oocytes expressing cysteine-substitution mutants (G514C and E518C) were incubated in 10 mM DTT-containing frog Ringer’s solution (88 mM NaCl, 1 mM KCl, 2.4 mM NaHCO_3_, and 0.3 mM Ca(NO_3_)_2_, 0.41 mM CaCl_2_, 0.82 mM MgSO_4_, and 15 mM HEPES-NaOH [pH 7.4] with 0.1% penicillin-streptomycin) at 20°C-25°C for 15 min, and were thoroughly washed with DTT-free Ringer’s solution. All measurements were performed at 20°C-25°C and conducted within 30 min so as to avoid the formation of nonspecific disulfide bonds (*Liu et al., 2002; Zhang et al., 2007*). The bath solution was ND-96 (96 mM NaCl, 2 mM KCl, 1.8 mM CaCl_2_,1 mM MgCl_2_, and 5 mM HEPES-NaOH [pH 7.4]). The holding membrane potential was set at −90 mV. For hERG *G*-*V* relationship experiments (WT and mutants except for L520A and L523A), currents were elicited by depolarizing voltage step pulses from −80 mV to +60 mV in 10-mV increments for 1 s, followed by a step pulse to −60 mV for 1 s. Due to the negatively shifted activation voltage of L520A and L523A, the magnitude of the depolarizing step pulses was individually adjusted for each of these mutants.

To determine the *K*_d_ value of APETx1 binding to hERG, the initial APETx1 concentration in the bath solution of 0.1 μM, and the concentration was increased cumulatively up to 10 μM. For mutational screening of hERG, APETx1 was added to the bath solution at a concentration of 10 μM, which is maximal concentration that oocytes can tolerate without leakage. The voltage-pulse protocol for *G*-*V* relationships was performed after the effect of APETx1 reached a steady state.

### Analysis of *G*-*V* relationships

All curve-fittings were performed using MATLAB R2019b (MathWorks Inc., Natick, MA). The *G*-*V* curves were fitted with the Boltzmann function:

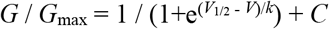

where the *G* / *G*_max_ is measured conductance relative to the maximal conductance, as determined from the peak of the outward tail current at −40 mV in TEVC recordings or at −60 mV in patch-clamp recordings; the *V*_1/2_ value is the membrane potential when the *G*-*V* relationship reaches half-maximal activation; the *k* value is the slope factor; and C is a constant component.

### Statistical analysis

All averaged data are presented as the mean ± SEM. The n value is the number of recordings. Multiple-group comparison was performed by one-way analysis of variance (ANOVA) followed by Tukey’s test using IBM SPSS Statistics, Version 25 (IBM Corp. Armonk, NY.). *p* < 0.05 was considered statistically significant (*,0.01 ≤ *p* < 0.05; **, 0.001 ≤ *p* < 0.01; ***, *p* < 0.001).

### Construction of the structural model of the APETx1-VSD complex

Homology models of the hERG VSD (residues 398-545) showing the S4-up or S4-down conformation were built using MODELLER 9.23 (Webb and Sali, 2016). The cryo-EM structure of rEAG1(*Whicher and MacKinnon, 2016*) was utilized as a template. The intact sequence alignment was used for the up model, whereas the three- and six-residue downward shift sequence alignments of S4 were used for the one- and two-helical-turn down models, respectively.

The structural model of the APETx1-hERG complex was generated with the HADDOCK2.4 web server (*van Zundert et al., 2016*). The unambiguous distance restraints were tabulated so that the key APETx1 residues (F15, Y32, F33, and L34) and the key hERG residues (F508, E518, and I521) would be located within 3.0 Å or less. The hydrogen bond restraints were also specified between the APETx1 Y32 and hERG E518 side chains. For rigid-body energy minimization, 1000 structures with the 200 lowest energy solutions were generated and used for subsequent semi-flexible simulated annealing and water refinement. Molecular graphics figures were depicted using CueMol (http://www.cuemol.org/).

## Acknowledgements

We are grateful to SB Drug Discovery Limited for providing SB-HEK-hERG. We also thank Ms. C. Naito of the laboratory of Y. Kubo for technical support. This research was supported in part by Japan Society for the Promotion of Science KAKENHI Grant Numbers JP17H03978 and JP19H04973 (to M.O.), a grant from The Vehicle Racing Commemorative Foundation (to M.O.), a grant from Takeda Science Foundation (to M.Y. and M.O.), and Platform Project for Supporting Drug Discovery and Life Science Research (Basis for Supporting Innovative Drug Discovery and Life Science Research (BINDS)) from AMED under Grant Numbers JP20am0101073 (support number 0928) and JP18am0101033 (support number 0004) (to Y.N.). We would like to thank Editage (www.editage.com) for English language editing.

