## Supplementary Figure for "Mechanism of hERG inhibition by gating-modifier toxin, APETx1, deduced by functional characterization"

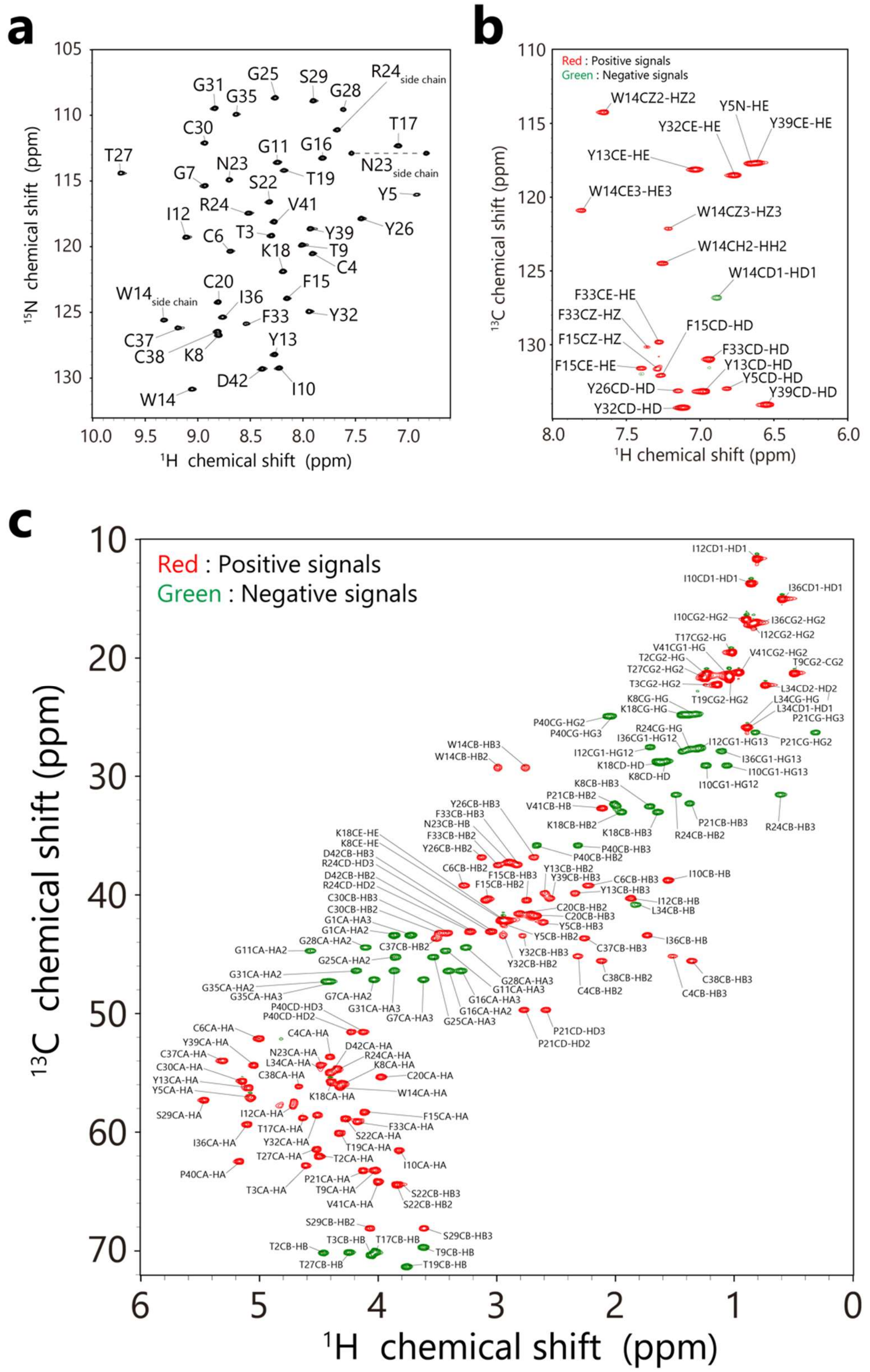


**Supplementary Figure 1. NMR resonance assignments of APETx1 on the $^{1}H$-$^{15}N$ HSQC and $^{1}H$-$^{13}C$ HSQC spectra observed at pH 6.0 and 298 K.**

$^{1}H$-$^{15}N$ HSQC spectrum (a), $^{1}H$-$^{13}C$ HSQC spectra for the aromatic region (b), and for the aliphatic region (c).

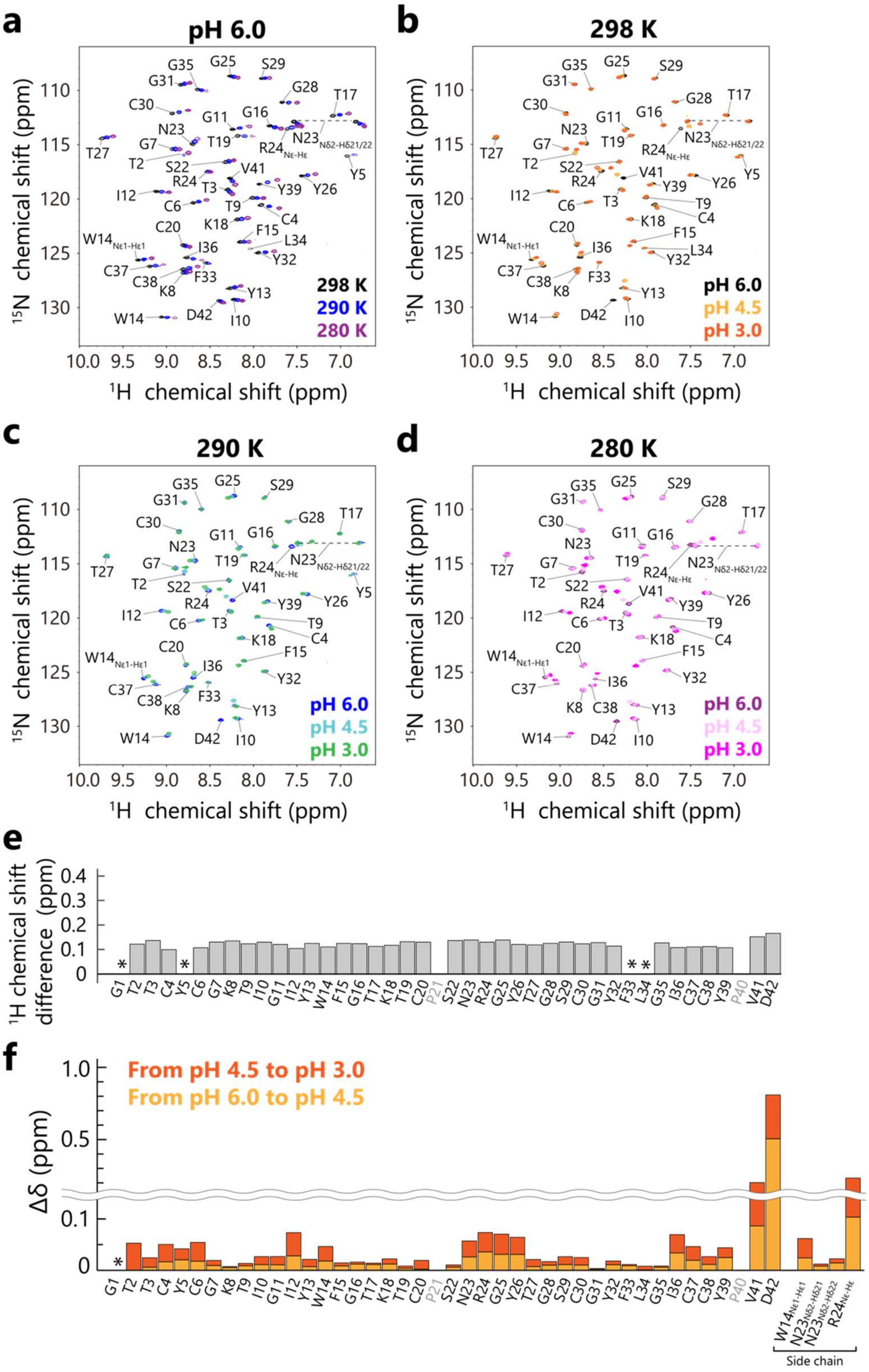


**Supplementary Figure 2. Variable-temperature and pH-titration NMR measurements using $^{1}$H-$^{15}$N HSQC.** (a) Overlay of the $^{1}$H-$^{15}$N HSQC spectra at pH 6.0 under the different temperature conditions. (b-d) Overlay of the $^{1}$H-$^{15}$N HSQC spectra at 298 K, 290 K, and 280 K in (b), (c), and (d), respectively, under different pH conditions. (e) $^{1}$H chemical shift differences of the backbone amide protons between recombinant APETx1 from the present study

(BMRB ID: 36345) and the natural product from a previous study (BMRB ID: 6370) (*Chagot et al., 2005*) under the same conditions (pH 3.0 and 280 K). The $^{1}H$ chemical shift values found in the previous study were subtracted from those found in the present study. The proline residues, which lack amide protons, are labeled in gray. Asterisks (*) show that the amide signal was not observed at pH 3.0 and 280 K. (f) pH-dependent chemical shift change (Δδ) values of the $^{1}H$-$^{15}N$ HSQC spectral signals at 298 K (b). The Δδ values were calculated using the following equation(*Mulder et al., 1999*):

$$\Delta\delta = \sqrt{[(\Delta\delta_{1H})^2 + (\Delta\delta_{15N} / 6.5)^2]}$$

where $\Delta\delta_{1H}$ and $\Delta\delta_{15N}$ are the chemical shift changes in the $^{1}H$ and $^{15}N$ dimensions, respectively. The proline residues, which lack amide protons, are labeled in gray. An asterisk (*) represents G1, which was not observed in the $^{1}H$-$^{15}N$ HSQC spectra.

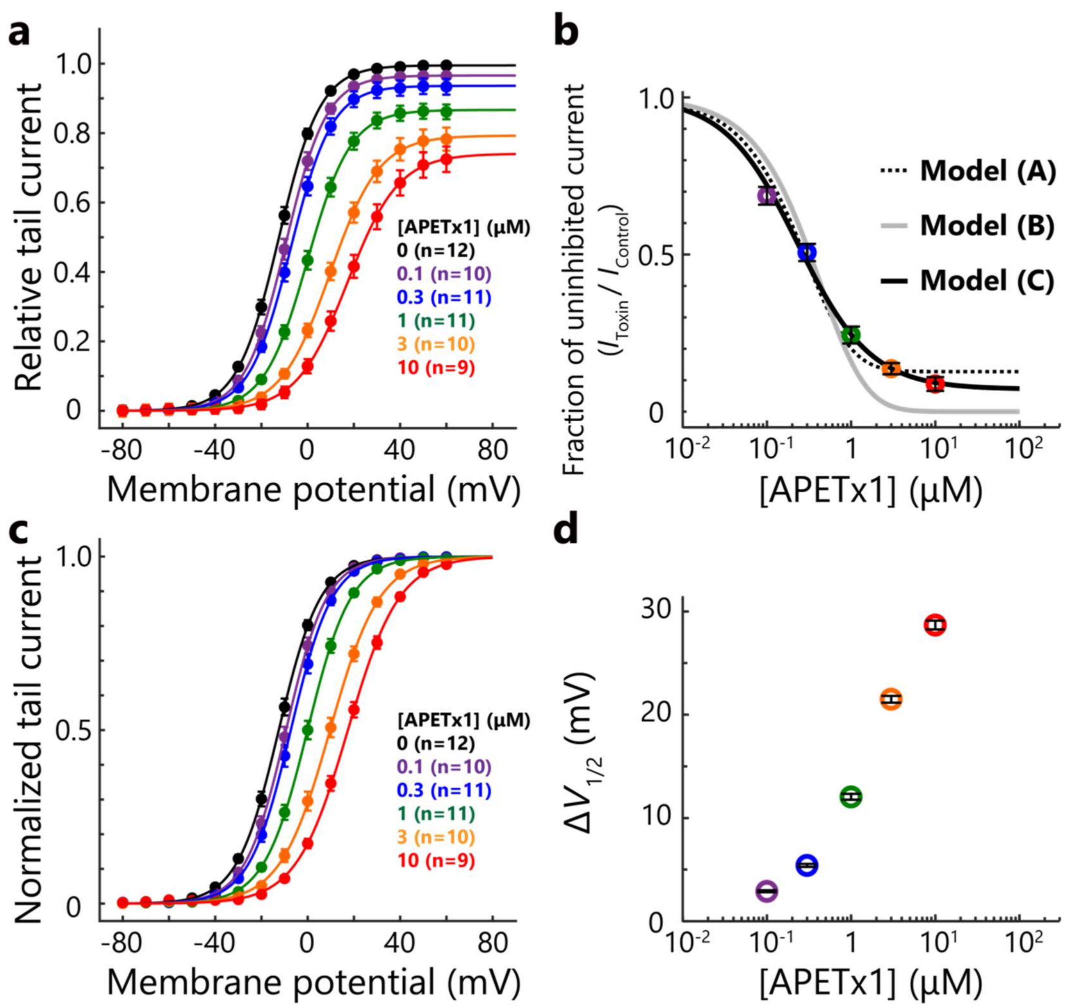


**Supplementary Figure 3. Dose-dependent effects of APETx1 on hERG in TEVC recordings.**

(a) *G-V* curves (mean ± SEM) of hERG in the presence or absence of different concentrations of APETx1. (b) Dose-response curve using the fraction of uninhibited currents at the depolarizing pulse, −30 mV. We fitted the data with the following three models (*Zhang et al., 2007*): (A) four equivalent and independent binding sites per channel with fractional toxin-sensitive current,

$I_{Toxin} / I_{control} = A_{max}(K_d / (K_d + [Tx]))^4 + (1 - A_{max})$, $K_d = 1.2$ μM, $A_{max} = 0.87$;

(B) Four equivalent and independent binding sites per channel with fully toxin-sensitive current,

$I_{Toxin} / I_{control} = (K_d / (K_d + [Tx]))^4$, $K_d = 1.7$ μM;

(C) One binding site per channel with fractional toxin-sensitive current,

$I_{Toxin} / I_{control} = A_{max} (K_d / (K_d + [Tx])) + (1 - A_{max})$, $K_d = 0.23$ μM, $A_{max} = 0.93$. (c) Normalized *G-V* curves (mean ± SEM). (d) The $\Delta V_{1/2}$ values of different concentrations of APETx1. Data points and error bars represent the mean values ± SEM.

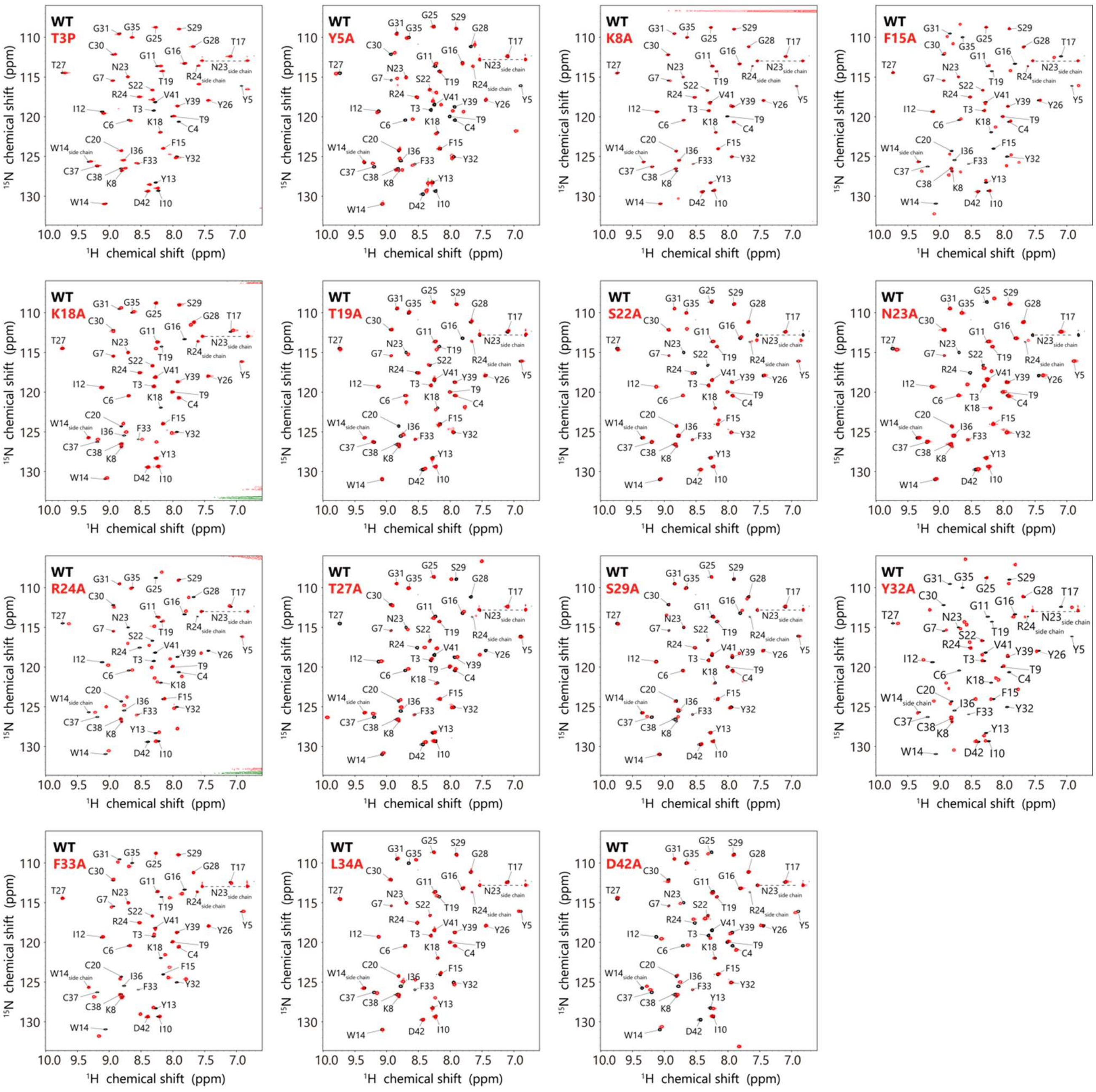


**Supplementary Figure 4. $^{1}H$-$^{15}N$ HSQC spectra of APETx1 mutants.**

All $^{1}H$-$^{15}N$ HSQC spectra were measured at pH 6.0 and 298 K in the following solution: T3P, K8A, F15A, K18A, R24A, Y32A, or F33A mutants, 20 mM potassium phosphate (pH 6.0), 100 mM KCl, and 10% $D_2O$; Y5A, T19A, S22A, N23A, T27A, S29A, L34A, or D42A mutants, 10% $D_2O$ (pH 3.0); and WT recorded in both conditions. The spectrum of each mutant is superimposed onto that of WT under identical solution conditions. NMR resonance assignments were labeled according to the signals of WT.

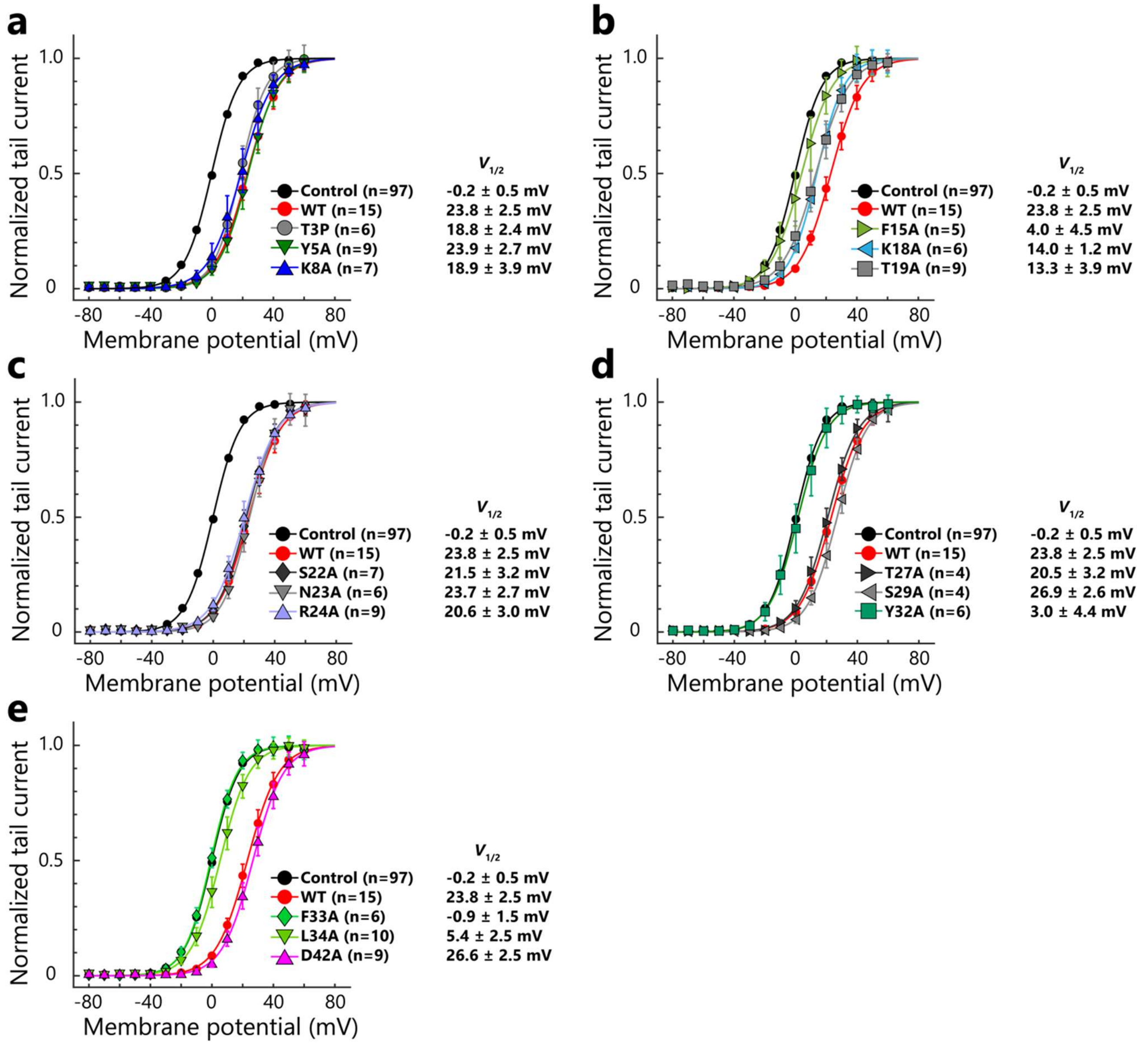


**Supplementary Figure 5. *G-V* curves of hERG in the presence or absence of 10 μM APETx1 and mutants.**
Normalized *G-V* curves (mean ± SEM) of hERG in the presence of 15 APETx1 mutants are sorted by amino acid residue order, divided by three into the five panels, (a) to (e).

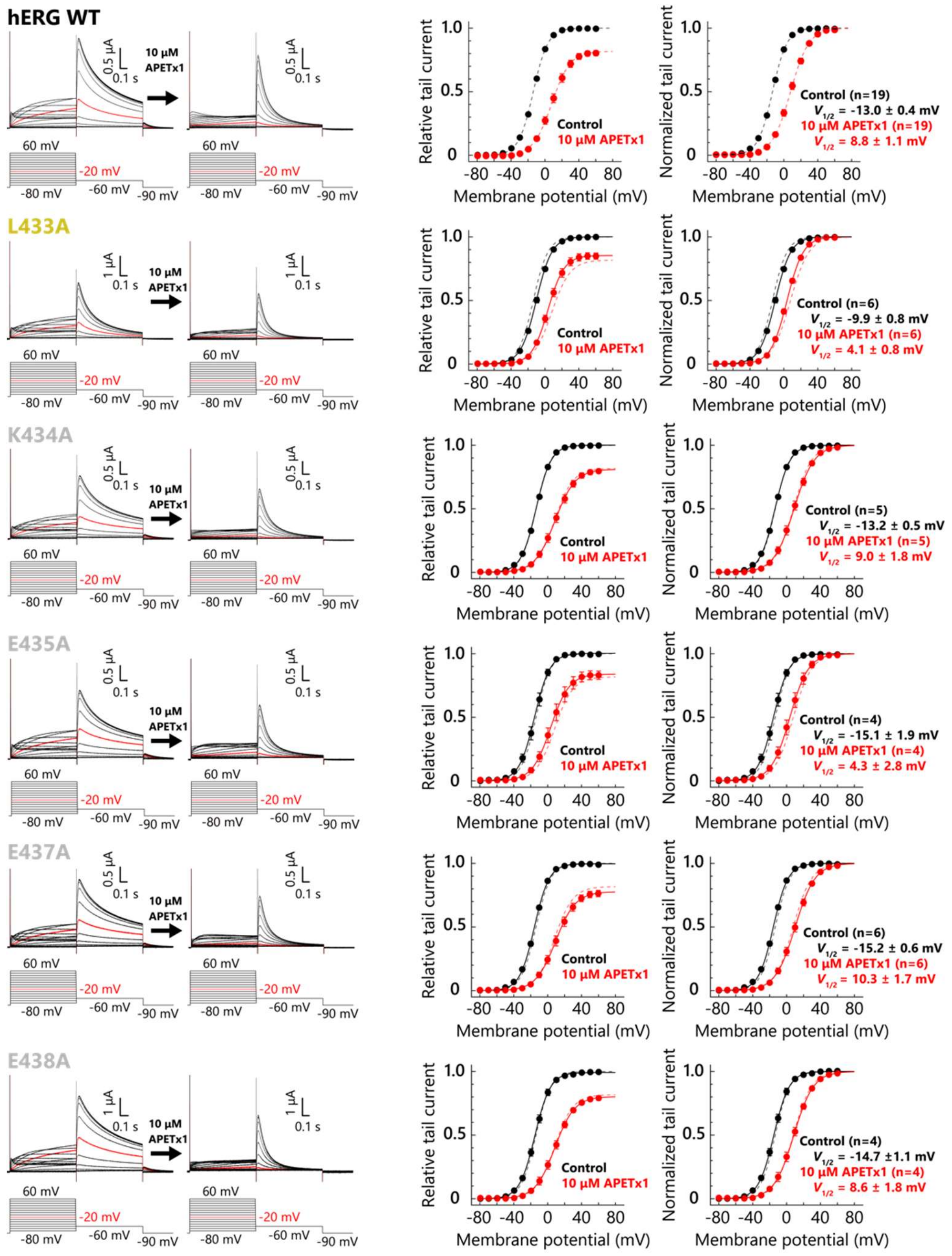


**Supplementary Figure 6. The current traces and *G-V* curves of hERG and its mutants in the presence or absence of 10 μM APETx1.**

Current traces of the hERG mutants before and after the administration of 10 μM APETx1 (left). Voltage protocol is illustrated at the bottom of each current trace. Current traces and voltage protocols at arbitrary potentials are depicted with red to clearly indicate the current reduction by the inhibitory effect of APETx1. G-V curves (mean ± SEM) of hERG mutants in the absence (black filled circle and solid line) or presence (red filled circle and solid line) of 10 μM APETx1 (right). The fitting curves of the WT in the absence (black dashed line) and presence (red dashed line) of 10 μM APETx1 are superimposed onto those of the mutants.

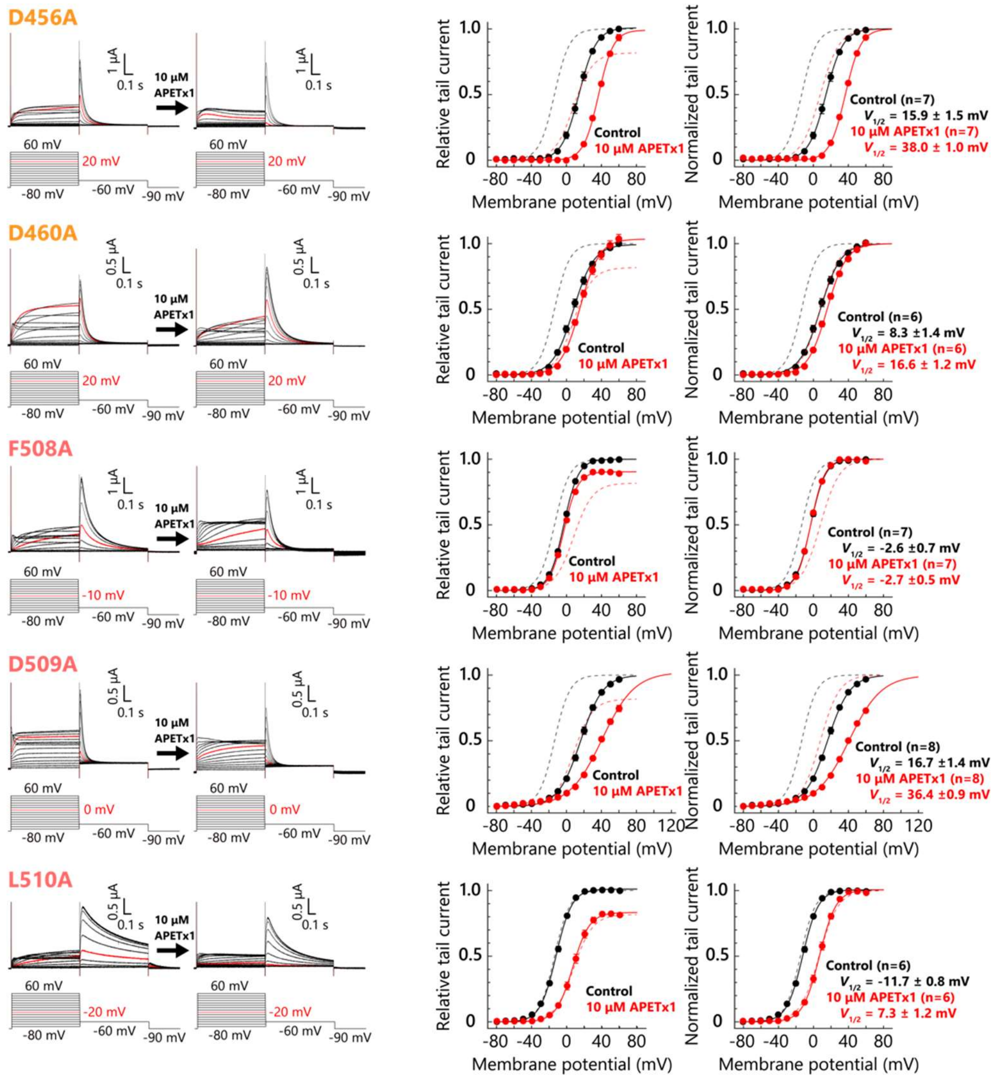


**Supplementary Figure 7. The current traces and *G-V* curves of hERG mutants in the presence or absence of 10 μM APETx1.**

Current traces of the hERG mutants before and after the administration of 10 μM APETx1 (left). Voltage protocol is illustrated at the bottom of each current trace. Current traces and voltage protocols at arbitrary potentials are depicted with red to clearly indicate the current reduction by the inhibitory effect of APETx1. G-V curves (mean ± SEM) of hERG mutants in the absence (black filled circle and solid line) or presence (red filled circle and solid line) of 10 μM APETx1 (right). The fitting curves of the WT in the absence (black dashed line) and presence (red dashed line) of 10 μM APETx1 are superimposed onto those of the mutants.

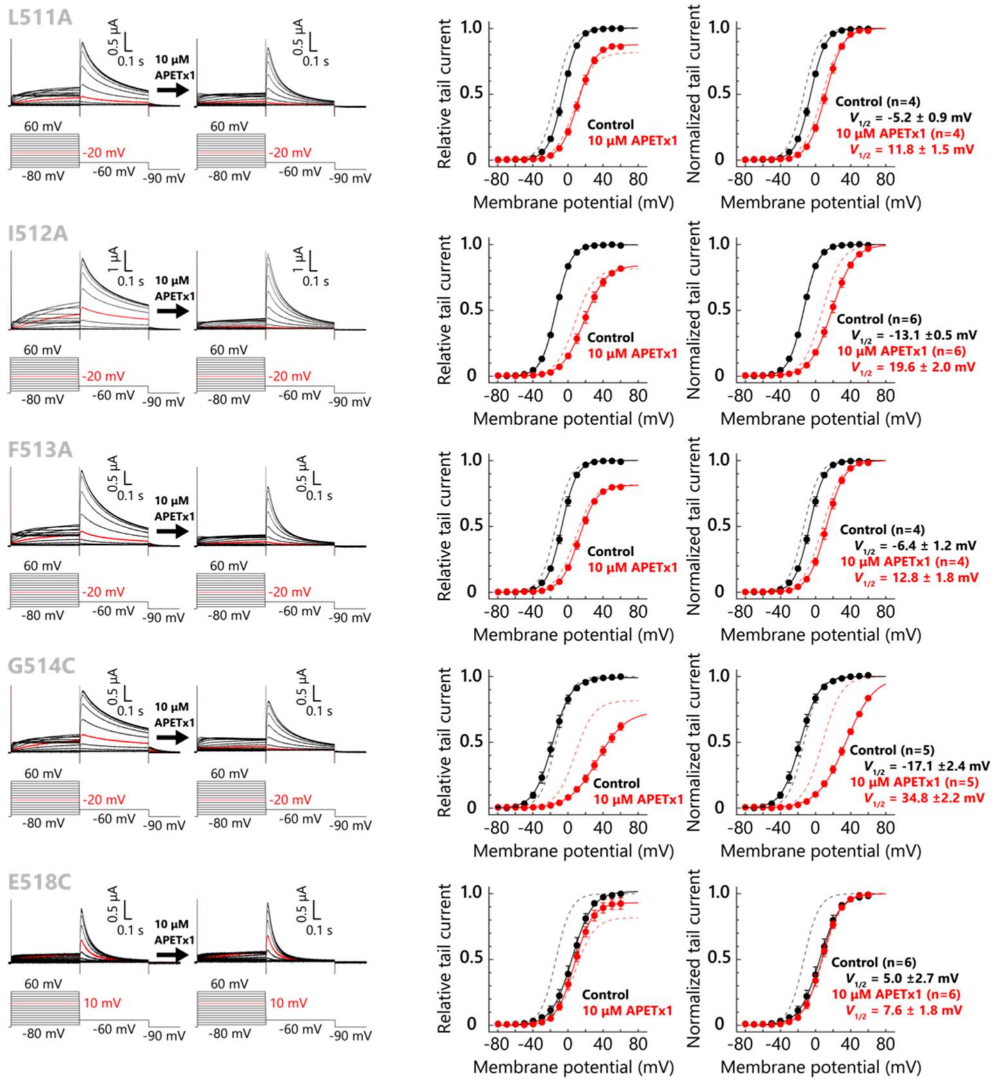


**Supplementary Figure 8. The current traces and *G-V* curves of hERG mutants in the presence or absence of 10 µM APETx1.**

Current traces of the hERG mutants before and after the administration of 10 µM APETx1 (left). Voltage protocol is illustrated at the bottom of each current trace. Current traces and voltage protocols at arbitrary potentials are depicted with red to clearly indicate the current reduction by the inhibitory effect of APETx1. G-V curves (mean ± SEM) of hERG mutants in the absence (black filled circle and solid line) or presence (red filled circle and solid line) of 10 µM APETx1 (right). The fitting curves of the WT in the absence (black dashed line) and presence (red dashed line) of 10 µM APETx1 are superimposed onto those of the mutants.

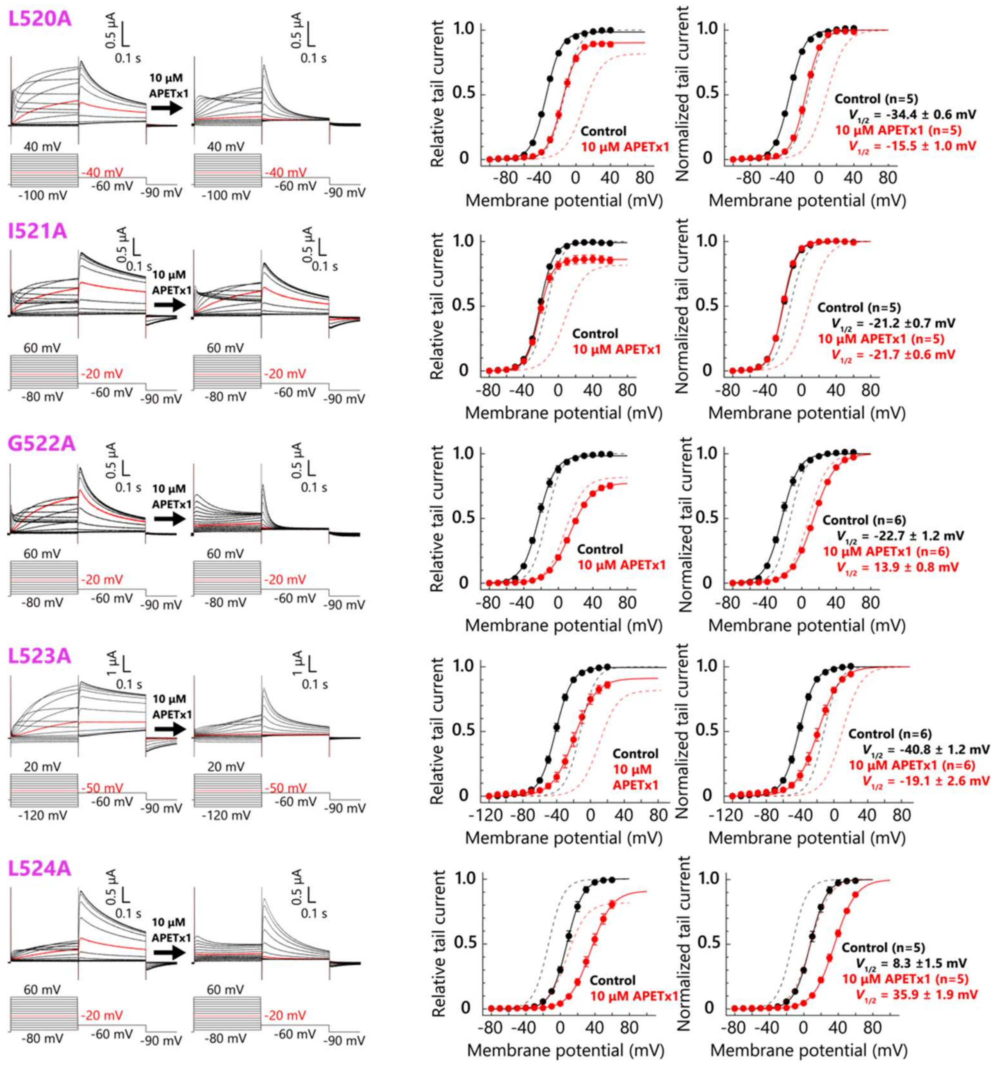


**Supplementary Figure 9. The current traces and *G-V* curves of hERG mutants in the presence or absence of 10 µM APETx1.**

Current traces of the hERG mutants before and after the administration of 10 µM APETx1 (left). Voltage protocol is illustrated at the bottom of each current trace. Current traces and voltage protocols at arbitrary potentials are depicted with red to clearly indicate the current reduction by the inhibitory effect of APETx1. G-V curves (mean ± SEM) of hERG mutants in the absence (black filled circle and solid line) or presence (red filled circle and solid line) of 10 µM APETx1 (right). The fitting curves of the WT in the absence (black dashed line) and presence (red dashed line) of 10 µM APETx1 are superimposed onto those of the mutants.

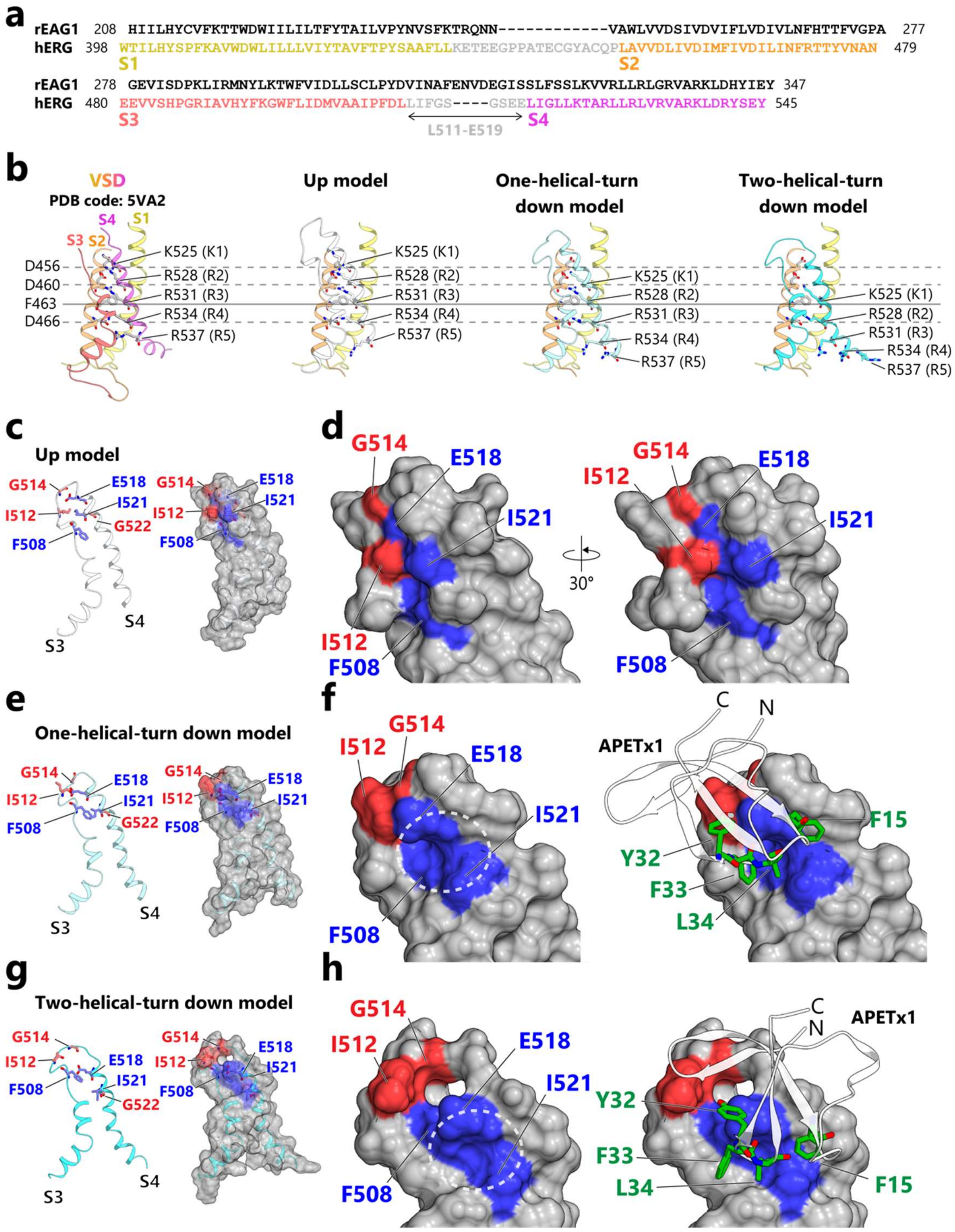


**Supplementary Figure 10. Model building of the APETx1-VSD complex.**

(a) Sequence alignment of the VSD of hERG (The Universal Protein Resource Knowledgebase (UniProtKB) Entry: Q12809) and rEAG1 (UniProtKB Entry: Q63472). The sequence of hERG is colored as in Figure 4a. (b) Validation of the homology models of the hERG S3-S4 region. Ribbon representation of the cryo-EM structure of hERG (PDB code: 5VA2; *Wang and MacKinnon, 2017*) up model, one- and two-helical-turn down models (from left to right). Basic residues at positions K1-R5 on S4, and the gating charge transfer center residue (F463) and the acidic residues

(D456, D460, and D466) on S2 are represented as sticks. To clearly show the position of S4, the Cα positions of the S2 residues are depicted with horizontal gray dashed (D456, D460, and D466) or solid lines (F463). (c) Ribbon and semi-transparent surface representations of the up model of the S3-S4 region, colored as in Figure 5. (d) Close-up view of (c). The one- (e) and two-helical-turn down models (g) of the S3-S4 region are represented as ribbons and semi-transparent surfaces colored as in Figure 5. On the left of (f) and (h) are close-up views of (e) and (g), respectively. White dotted circles indicate the crevice between F508, E518, and I521. On the right, docked APETx1 is also displayed as semi-transparent ribbons with the green sticks indicating key residues involved in hERG inhibition.

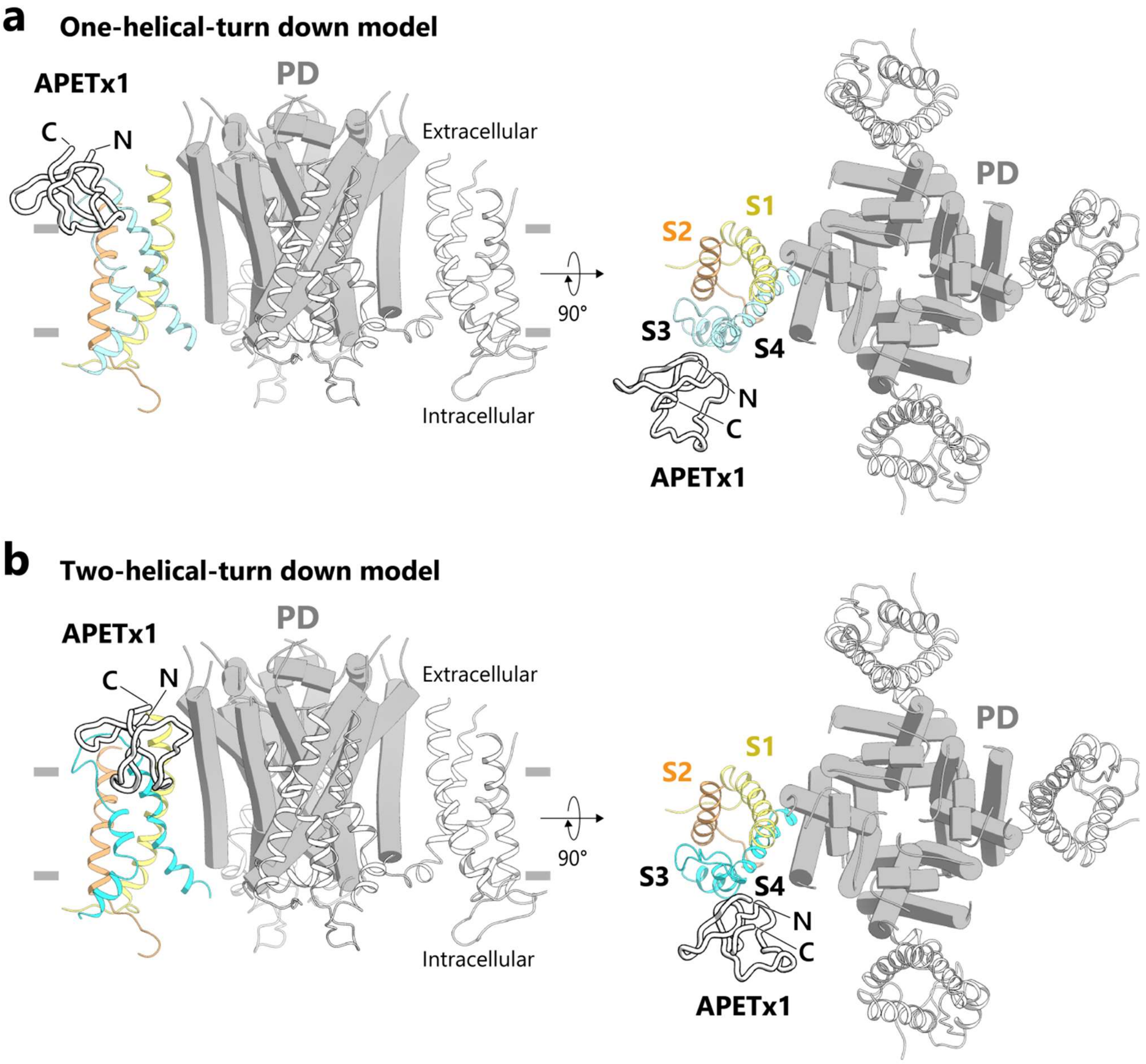


**Supplementary Figure 11. The binding location of APETx1 in a tetrameric transmembrane architecture of hERG.**

APETx1-VSD complex models, in which S4 adopts one- and two-helical-turn down conformations in (a) and (b), respectively, are superimposed onto the transmembrane domain of the cryo-EM of hERG (PDB code: 5VA2; *Wang and MacKinnon, 2017*), viewed from within the membrane plane (left) and from the extracellular side (right). One of the four VSDs of the cryo-EM structure is substituted by the APETx1-VSD complex model. The PD is represented as a cartoon, the VSDs as ribbons, and APETx1 as tubes. In the APETx1-docked VSD, S1 is yellow; S2, orange; S3 and S4 in (a), sky blue; and S3 and S4 in (b), cyan.
